# Temperature-dependent small RNA expression depends on wild genetic backgrounds of *Caenorhabditis briggsae*

**DOI:** 10.1101/2022.05.23.493161

**Authors:** Daniel D. Fusca, Eesha Sharma, Jörg G. Weiss, Julie M. Claycomb, Asher D. Cutter

## Abstract

Geographically distinct populations can adapt to the temperature conditions of their local environment, leading to temperature-dependent fitness differences between populations. Consistent with local adaptation, phylogeographically distinct *Caenorhabditis briggsae* nematodes show distinct fitness responses to temperature. The genetic mechanisms underlying local adaptation, however, remain unresolved. To investigate the potential role of small noncoding RNAs in genotype-specific responses to temperature, we quantified small RNA expression using high-throughput sequencing of *C. briggsae* nematodes from tropical and temperate strain genotypes reared under three temperature conditions (14°C, 20°C, 30°C). Strains representing both tropical and temperate regions showed significantly lower expression of PIWI-interacting RNAs (piRNAs) at high temperatures, primarily mapping to a large ∼7 Mb long piRNA cluster on chromosome IV. We also documented decreased expression of 22G-RNAs antisense to protein-coding genes and other genomic features at high rearing temperatures for the thermally-intolerant temperate strain genotype, but not for the tropical strain genotype. Reduced 22G-RNA expression was widespread along chromosomes and among feature types, indicative of a genome-wide response. Targets of the EGO-1/CSR-1 22G-RNA pathway were most strongly impacted compared to other 22G-RNA pathways, implicating the CSR-1 Argonaute and its RNA-dependent RNA polymerase EGO-1 in the genotype-dependent modulation of *C. briggsae* 22G-RNAs under chronic thermal stress. Our work suggests that gene regulation via small RNAs may be an important contributor to the evolution of local adaptations.

## Introduction

Temperature is a universal property of all ecosystems that influences diverse aspects of organismal fitness. In species that inhabit distinct temperature regimes across their geographic range, natural selection can lead different populations to evolve adaptations to the temperature conditions of their local habitat, creating within-species variation in the response to temperature stress. For example, populations of *Drosophila melanogaster* from tropical regions are more heat-tolerant than populations from cooler temperate regions (Rohmer et al. 2004). Similarly, populations of *Arabidopsis thaliana* found at high latitudes are more cold-tolerant than populations from low latitudes (Zhen and Ungerer 2008). Understanding the process of adaptation is a central theme of evolutionary biology, and so there is much interest in identifying the genetic basis of such local adaptations in diverse taxa (Ahrens et al. 2018). However, it is not clear in all cases what the genetic variants and molecular mechanisms underlying these local adaptations to temperature are, or how this variation contributes to differential fitness.

An excellent system for interrogating the molecular mechanisms of local adaptation is the nematode roundworm *Caenorhabditis briggsae*, a self-fertilizing hermaphrodite and close relative of the model organism *Caenorhabditis elegans*. Like *C. elegans*, *C. briggsae* has a widespread global range, and therefore has the potential to demonstrate local adaptation to varying habitats (Kiontke et al. 2011; Cutter 2015). In support of this possibility, *C. briggsae* isolates found at tropical and temperate latitudes form genetically distinguishable phylogeographic groups (Cutter et al. 2006; Félix et al. 2013). Examining how populations of *C. briggsae* found at different latitudes may show local adaptation, Prasad et al. (2011) demonstrated that strains of *C. briggsae* from 3 phylogeographic groups reared at varying temperatures showed distinct reproductive fitness responses. Strains from tropical latitudes had higher lifetime fecundity than temperate strains at high temperatures (30°C) but lower fecundity than temperate strains at low temperatures (14°C), consistent with local adaptation. Subsequent studies have further characterized how *C. briggsae* strains differ in their response to temperature, such as how strains differ genetically in behavioral responses to environmental temperature, including thermal preferences, isothermal tracking, motility, sperm and oocyte fertility, and acute cold sensitivity (Stegeman at al. 2013; Poullet et al. 2015; Stegeman et al. 2019; Wang et al. 2021). mRNA sequencing of a tropical and a temperate strain reared at extreme temperatures found a significant genotype-temperature interaction for 56% of the differentially-expressed genes (i.e. strains differed in temperature-dependent expression changes), many of which were associated with known biological processes such as translation and oogenesis (Mark et al. 2019). Many genes are upregulated in response to extreme cold stress in temperate, but not tropical, *C. briggsae* strains, and the role of a few of these genes in mediating strain-specific cold tolerance has been experimentally confirmed (Wang et al. 2021). Despite this work, the genetic mechanisms underlying these local adaptations in *C. briggsae* remain incompletely understood.

One potential contributor to locally-adaptive evolutionary responses that remains largely uncharacterized are small noncoding RNA regulatory pathways. Small RNAs (noncoding RNAs generally between 20 to 30 bases in length) associate with Argonaute proteins to regulate the expression of target loci, such as protein-coding genes and transposable elements, and include microRNAs, PIWI-interacting RNAs (piRNAs), and endogenous small interfering RNAs (endo-siRNAs) (Hoogstrate et al. 2014). Small RNA pathways can show adaptive differences between species (Palmer et al. 2018; Franchini et al. 2019), but the role of small RNA-mediated gene regulation in shaping differences between populations within a single species is less clear.

However, there is some evidence linking small RNAs to local adaptations. Comparisons between different populations have revealed microRNA loci in nematodes (Jovelin and Cutter 2011), humans (Torruella-Loran et al. 2016), and plants (Xie et al. 2017) that show strong allele frequency differentiation that is consistent with local adaptation, though the functional consequences of this differentiation are not fully understood. The genes that carry out small RNA regulation may also show population-specific signatures of positive selection, as seen for genes involved in the *D. melanogaster* piRNA pathway (Simkin et al. 2013). Sequence changes to small RNA binding sites can be involved in local adaptations as well, as has been reported for microRNA binding sites in populations of humans (Li et al. 2012) and *D. melanogaster* (Catalán et al. 2016). In addition to DNA sequence differences, natural populations can show variation in the expression of piRNAs (Ellison and Cao 2020), microRNAs (Arif et al. 2013), and siRNAs (Zhai et al. 2008). This variable expression between populations can cause intraspecific variation in morphology (Arif et al. 2013) or epigenetic regulation (Zhai et al. 2008), but the adaptive significance of this variation is usually uncharacterized. However, a few studies have identified differential microRNA expression between natural populations with known local adaptations, such as to heat stress (Graham and Barreto 2019) or hydrogen sulphide toxicity (Kelley et al. 2021), providing a link between microRNA-mediated gene regulation and locally-adaptive phenotypes. While much is unknown about how endo-siRNAs may be involved in local adaptation compared to microRNAs, the regulatory role of some endo-siRNAs in mediating responses to environmental stress (Wu et al. 2020) suggests that environmental differences between populations could lead to adaptive evolution in endo-siRNA pathways.

Studies in *C. elegans* indicate that small RNAs may play an important role in how these nematodes respond to temperature stress. For example, loss of 26G-RNAs (endo-siRNAs that are 26 bases long and begin with a 5’ guanine nucleotide) induces spermatogenesis defects at elevated temperatures (Conine et al. 2010). Additionally, production of piRNAs is repressed in response to increased temperatures (Belicard et al. 2018), and disruption of the piRNA pathway in *C. elegans* leads to fertility defects at high temperatures (Wang and Reinke 2008). It has been shown that some *C. briggsae* microRNAs have a high level of polymorphism between strains, and this variation may have functional consequences (Jovelin and Cutter 2011). Thus, differences in small RNA sequence and expression between tropical and temperate strains have the potential to contribute to the temperature-dependent mRNA transcriptome profiles and fitness differences observed in *C. briggsae*, although to date this idea has not yet been tested.

To explore this possibility, we characterize differential small RNA expression in response to chronic temperature stress for distinct temperate and tropical genotypes of *C. briggsae*. We observe genome-wide reductions in small RNA expression under high temperature stress that are specific to the temperate genotype, consistent with the reduced fitness of this strain at high temperatures. By classifying genes that are regulated by different small RNA pathways, we test how specific pathways may be responsible for this genotype-specific response. The temperature-dependent responses of distinct genotypes that we observe point to a role for the regulation of gene expression via small RNAs in the process of local adaptation.

## Results

### *C. briggsae* genotypes show contrasting patterns of microRNA expression at low temperatures

We investigated the effect of chronic temperature stress from rearing at 14°C, 20°C, and 30°C on the small RNAs produced by different *C. briggsae* genotypes, using two strains that represent tropical (AF16) and temperate (HK104) latitude phylogeographic groups of this species. Endo-siRNAs with a 5’ guanine and 22-nucleotide length (22G-RNAs) dominate regulatory small RNAs in these samples (Fig. 1), which is consistent with other *Caenorhabditis* nematodes (Hoogstrate et al. 2014). In total, 31.4% of small RNAs corresponded to 22G-RNAs, accompanied by major contributions of microRNAs (27.3%) and 21U-piRNAs (12.0%), as well as a minor contribution of 26G-RNAs (1.4%) (Fig. 1, Fig. S1). We found that 21.2% of all small RNAs aligned antisense to annotated protein-coding genes, with only 5.3% aligning in the sense orientation (Fig. S1), as expected for endo-siRNAs such as 22G-RNAs. About one quarter (25.3%) of small RNAs aligned to regions of the genome without any functional annotation, though this is likely due to unannotated loci not present in our *C. briggsae* genome annotation (e.g., piRNA loci). Small RNAs with a length of 24 nucleotides were strongly enriched for those starting with a 5’ cytosine (C) (Fig. 1), as has been seen previously in *C. elegans* and *C. briggsae* (Vasale et al. 2010; Tu et al. 2015, Charlesworth et al. 2021). This enrichment was due to a single highly-expressed microRNA, *cbr-mir-52* (WBGene00250149), located on chromosome IV. The strongest changes in microRNA expression were observed in temperate HK104 worms reared at 14°C, which showed a 1.7-fold reduction in overall microRNA expression relative to HK104 worms at 20°C (Fig. S1). Interestingly, the tropical AF16 genotype showed the opposite pattern, with overall microRNA expression being 1.3-fold higher at 14°C relative to 20°C (Fig. S1), suggesting possible genotype-specific differences in microRNA-mediated gene regulation at low temperatures.

**Figure 1.**
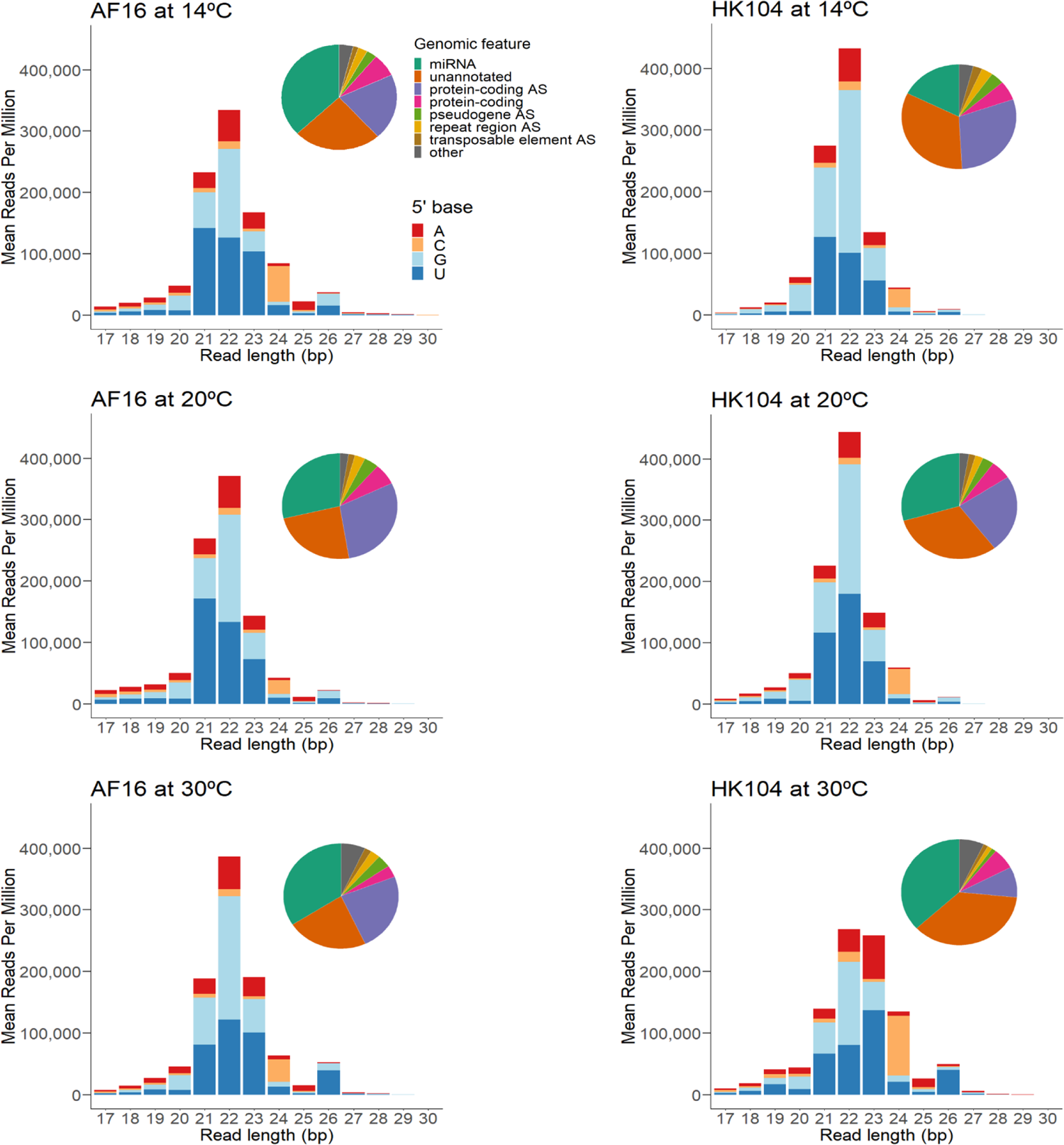
Mean expression of small RNAs of different lengths that mapped perfectly to the *C. briggsae* genome for each genotype-temperature combination, separated by the 5’ nucleotide of mapped small RNAs. Pie charts give the distribution of genomic features that small RNAs mapped to. “AS” indicates reads mapping antisense to a feature.

### Genomic clusters of 21U-piRNAs show distinct temperature-dependent expression

Expression of *C. briggsae* piRNAs (which are 21 bases long and begin with a 5’ uracil nucleotide) originated primarily from three major clusters, two encoded on chromosome IV and one on chromosome I (Fig. 2), in line with previous studies (Shi et al. 2013, Beltran et al. 2019). We found that, overall, 21U-piRNAs show lower expression in worms reared at high temperature (30°C) for both the tropical (AF16) and temperate (HK104) genotypes (Fig. 1, Fig. 2). This trend was driven almost entirely, however, by 21U-piRNAs that derived from the large piRNA gene cluster on the left arm of chromosome IV (0.6 Mbp to 7.0 Mbp) (Fig. 2). The smaller piRNA cluster on the right arm of chromosome IV (13.2 Mbp to 15.0 Mbp), by contrast, showed a reciprocal pattern: elevated 21U-piRNA expression at low temperature (14°C) (Fig. 2). piRNAs aligning to chromosome I showed yet a third pattern: increased expression at high temperatures (most strongly in HK104, Fig. 2), although this was due to loci outside of the main piRNA cluster on this chromosome in addition to loci within the cluster (9.8 Mbp to 11.3 Mbp; Fig. S2). These observations point to the possibility of distinct, coordinated functions for those piRNAs encoded in different genomically-localized piRNA clusters in response to environmental or other perturbations.

**Figure 2.**
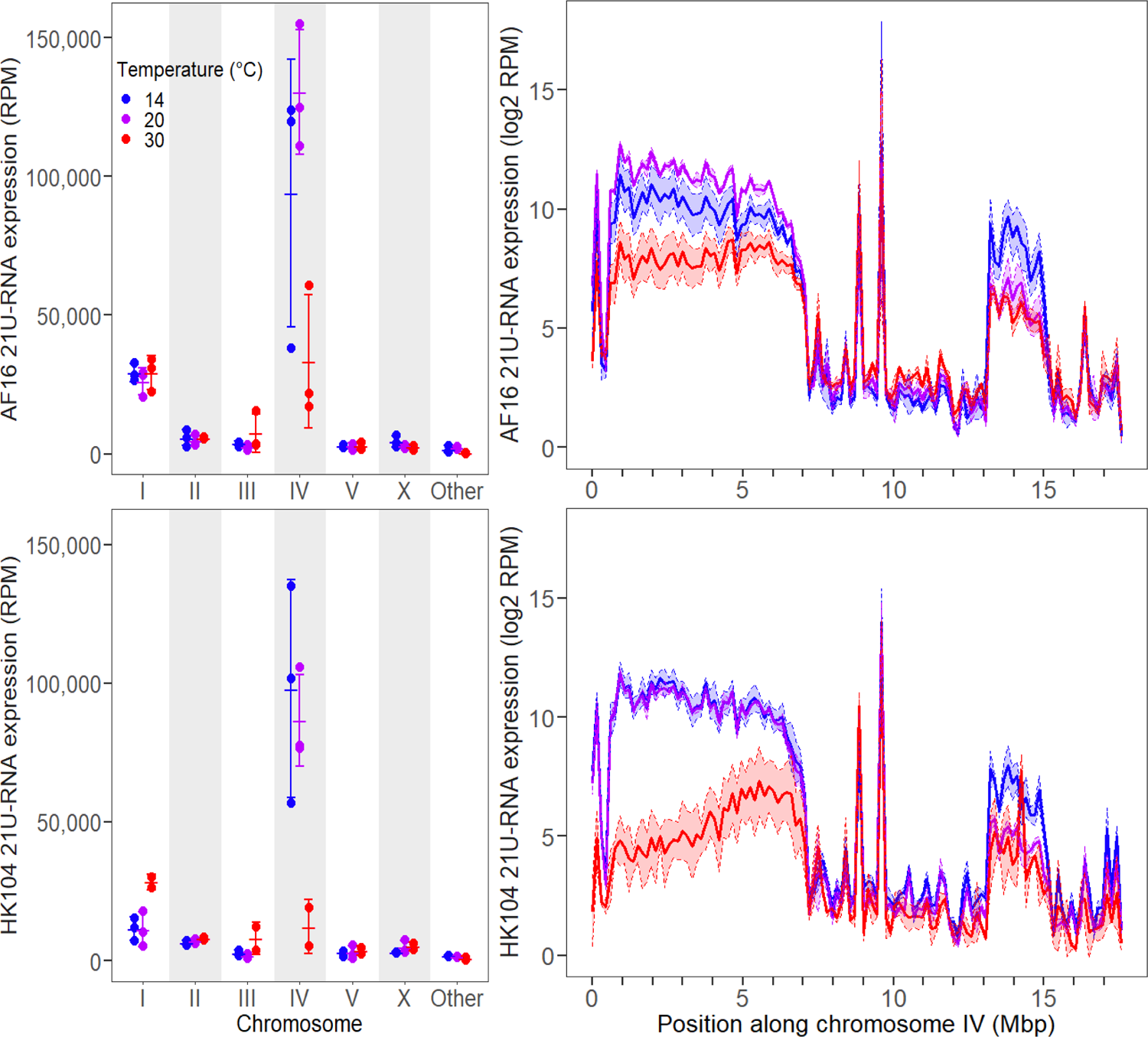
Total expression of 21U-RNAs aligning to each chromosome (left), as well as along the length of chromosome IV using genomic intervals 150kb wide (right), in AF16 (top) and HK104 (bottom) worms. Expression is in units of Reads Per Million (RPM) (left) or log2 Reads Per Million (right). Error bars and shaded regions give ±1 standard deviation around the mean for each set of replicates. “Other” refers to genomic scaffolds of unknown chromosomal origin.

### High temperatures reduce 22G-RNA expression only in the temperate genotype

In addition to identifying some shared temperature-dependent expression responses, we also observed genotype-specific small RNA expression changes in response to temperature. Using multidimensional scaling, expression profiles of 22G-RNAs that mapped antisense to annotated genomic features clustered into three distinct sample types: AF16 animals from all temperatures, HK104 worms from 14°C and 20°C, and a separate group of HK104 reared at 30°C (Fig. S3). Using linear modeling to compare expression between genotype-by-temperature combinations, temperate HK104 animals raised at 30°C showed a 2.3-fold decrease in 22G-RNA reads aligning antisense to protein-coding genes relative to HK104 animals raised at 20°C (*p* = 0.014; 22G-RNAs target protein-coding genes through sequence complementarity (Hoogstrate et al. 2014)), despite tropical AF16 animals having on average nearly identical gene-antisense 22G-RNA expression between 20°C and 30°C (*p* = 0.956; Fig. 1, Fig. 3, Fig. S1). This represented a significant genotype-temperature interaction on 22G-RNA expression at 30°C (*p* = 0.050), consistent with especially strong perturbation of 22G-RNA expression in HK104 at high temperatures. This trend also held for 22G-RNAs aligning antisense to pseudogenes (2.3-fold lower in HK104 at 30°C; *p* = 0.016) (Fig. 3), and was nominally similar if non-significant for repeat regions (1.6-fold lower; *p* = 0.338) and transposable elements (1.4-fold lower; *p* = 0.345), indicative of a genome-wide rather than locus-specific reduction in 22G-RNA expression for the temperate HK104 genotype reared at 30°C.

**Figure 3.**
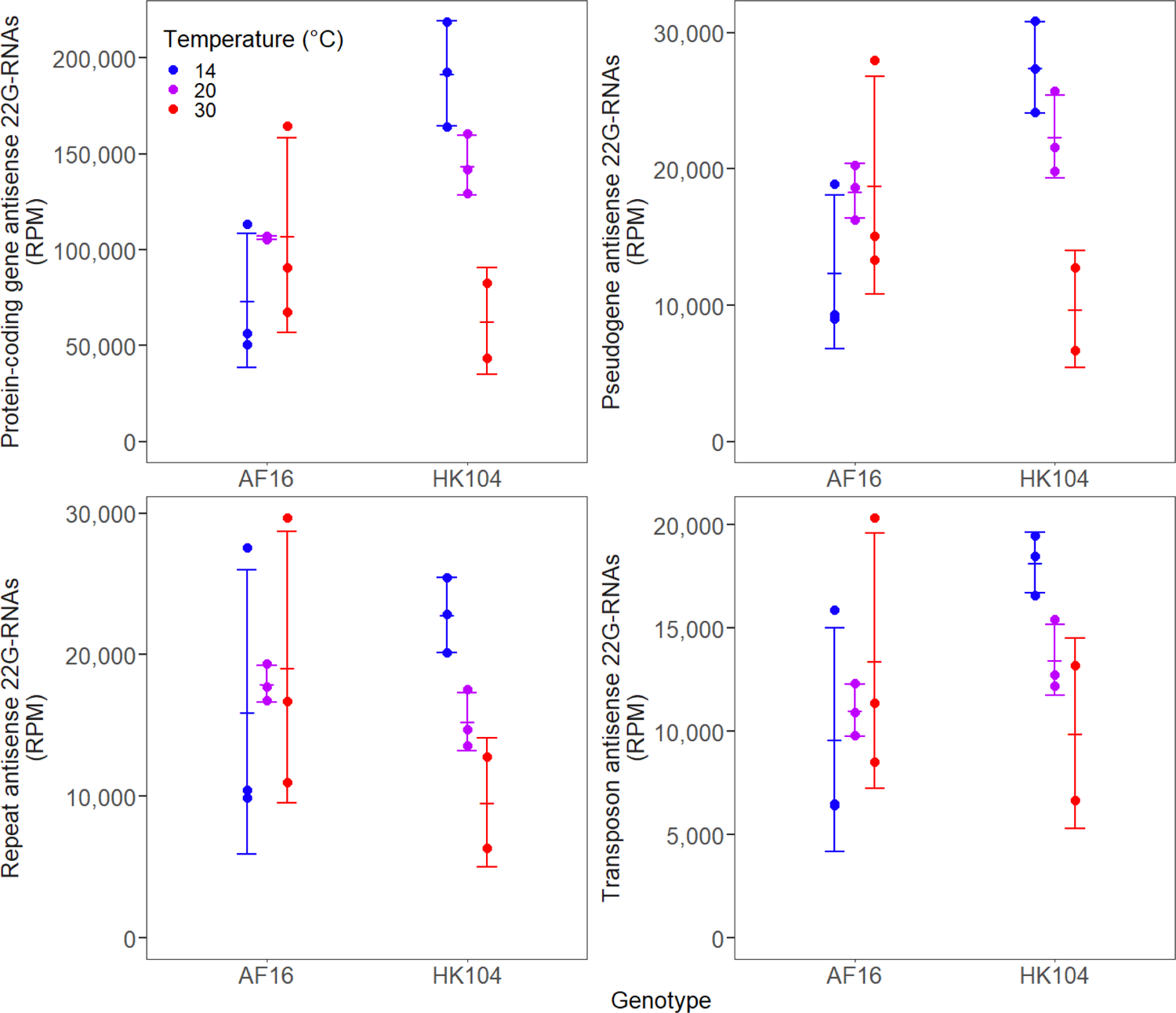
Total expression of 22G-RNAs aligning antisense to protein-coding genes, pseudogenes, repeat regions, and transposons for each genotype-temperature combination, considering only the genes, pseudogenes, repeat classes, and transposon classes that are present in the genome annotations for both strains. Expression is in units of Reads Per Million (RPM). Error bars give ±1 standard deviation around the mean of each set of replicates.

Supporting the idea of a global regulatory difference, we observed that expression of all 22G-RNAs across all six *C. briggsae* chromosomes decreased 1.5-fold for HK104 animals reared at 30°C compared to 20°C, with decreases ranging from 1.3-fold on chromosome III to 2.0-fold on chromosome II (Fig. 4, Fig. S4). Linear modeling of the 22G-RNAs aligning to each chromosome again found that, accounting for differences between chromosomes, this genome-wide decrease at 30°C is statistically significant (*p* = 0.0013). Tropical AF16 animals, by contrast, showed 1.2-fold higher average 22G-RNA expression at 30°C than at 20°C on all chromosomes (ranging from 1.03-fold higher on chromosome I to 1.3-fold higher on chromosome IV) except chromosome II, which had 1.1-fold lower average expression (Fig. 4), although this overall increase at 30°C was not statistically significant (*p* = 0.161). These contrasting patterns between genotypes again resulted in a significant genotype-temperature interaction at 30°C (*p* = 9.3 x 10^-4^). 22G-RNA expression was significantly higher on the X chromosome compared to all other chromosomes, regardless of genotype or temperature treatment (*p <* 6.1 x 10^-8^; Fig. 4).

**Figure 4.**
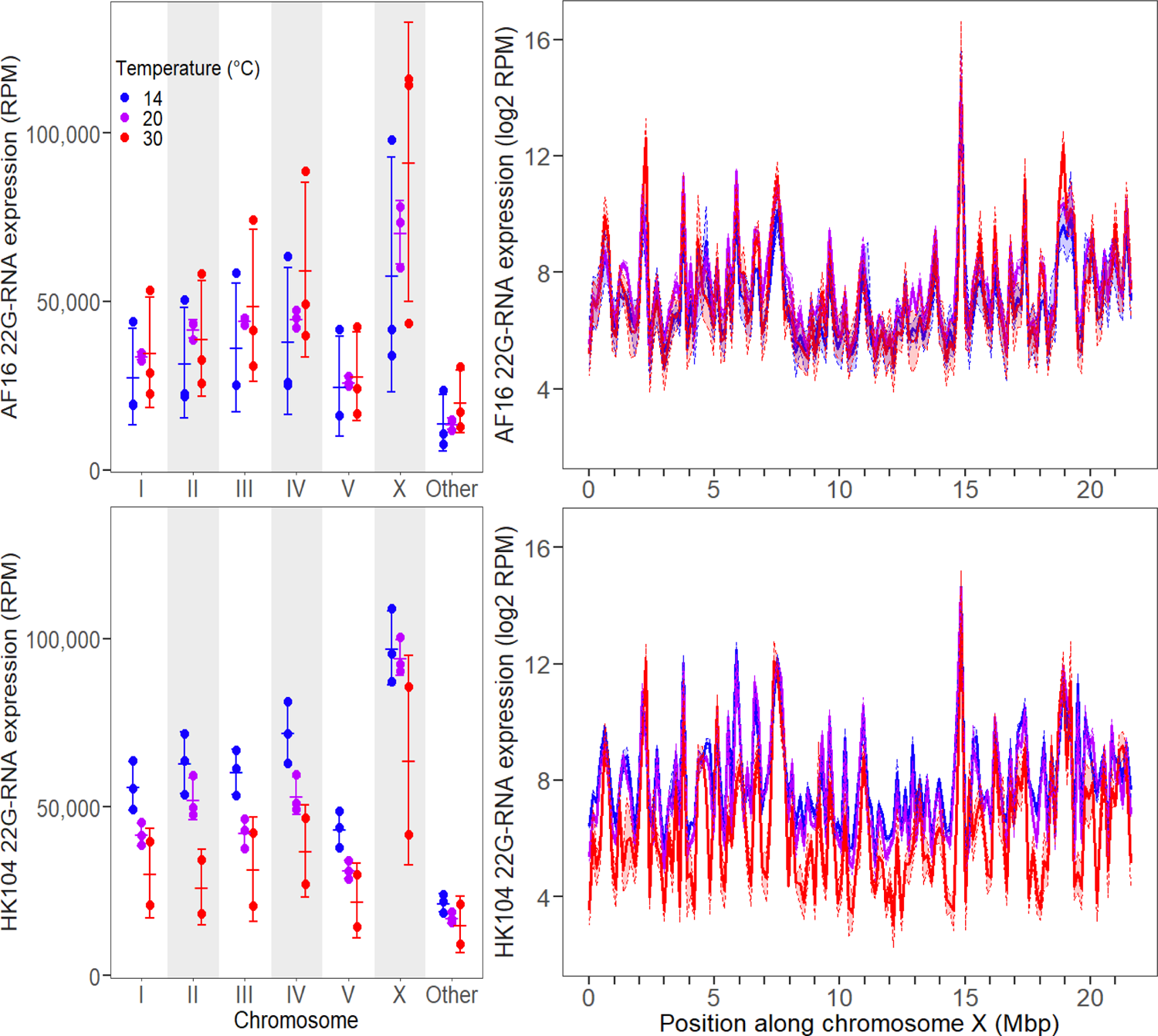
Total expression of 22G-RNAs aligning to each chromosome (left), as well as along the length of the X chromosome using genomic intervals 150kb wide (right), in AF16 (top) and HK104 (bottom) worms. Expression is in units of Reads Per Million (RPM) (left) or log2 Reads Per Million (right). Error bars and shaded regions give ±1 standard deviation around the mean for each set of replicates. “Other” refers to genomic scaffolds of unknown chromosomal origin.

Dividing the genome into 100 kb bins, 82% to 90% of bins across HK104 chromosomes showed this pattern of depressed 22G-RNA expression at 30°C versus just 52% to 78% of bins for AF16 (Table 1). Moreover, the mean magnitude of decreased expression was larger for HK104 than AF16 on all 6 chromosomes (Table 1). Both the chromosome arm and center domains reflected these patterns (Table S1, Table S2), although centers showed a more pronounced difference between genotypes: at 30°C, 35.4% of bins on chromosome centers had lower 22G-RNA expression in HK104 but not AF16, compared to only 19.5% of bins on chromosome arms (Fisher’s exact test, *p* = 9.6 x 10^-9^).

**Table 1:** Proportion of 100kb bins along each chromosome where mean 22G-RNA expression was lower at 30°C than it was at 20°C, for both genotypes. Bins showing decreased expression at 30°C were used to calculate the mean magnitude of the expression decrease for both genotypes. Expression decreases are in units of log_2_ Reads Per Million.

| Chromosome | % of bins in AF16<br>with decreased<br>expression at 30 °C | % of bins in HK104<br>with decreased<br>expression at 30 °C | Average expression<br>decrease for AF16<br>bins (log<sub>2</sub> RPM) | Average expression<br>decrease for HK104<br>bins (log<sub>2</sub> RPM) |
| --- | --- | --- | --- | --- |
| I | 64% (100 / 156) | 88% (137 / 156) | 0.51 | 1.40 |
| II | 64% (107 / 167) | 90% (150 / 167) | 0.61 | 1.53 |
| III | 52% (77 / 147) | 88% (129 / 147) | 0.49 | 1.37 |
| IV | 64% (113 / 176) | 82% (145 / 176) | 0.62 | 1.35 |
| V | 63% (123 / 196) | 82% (160 / 196) | 0.56 | 1.18 |
| X | 78% (169 / 217) | 85% (184 / 217) | 0.72 | 1.84 |

Using the 22G-RNA expression profiles of 14,283 genomic features to which 22G-RNA sequence reads aligned with antisense orientation (including protein-coding genes, pseudogenes, repeats, and transposable elements), we detected significant differential expression for 8531 of them due to strain genotype, rearing temperature, or a non-additive genotype-temperature interaction (i.e., expression changes due to temperature that differed between the AF16 and HK104 genotypes) (Fig. S5; Table S3). Most commonly, 57.7% of the features with differentially expressed 22G-RNAs showed genotype-temperature interactions (4924 features). Additionally, 1445 features (16.9%) showed expression differences due to strain genotype alone, 1415 (16.6%) due to temperature alone, and 747 (8.8%) due to a simple additive combination of both genotype and temperature. Of the 4924 features showing a genotype-temperature interaction, 64.0% (3151 features) recapitulated the distinctive overall reduction of 22G-RNAs in HK104 reared at 30°C relative to 20°C (Fig. S5). Compared to the genome-wide distribution of all protein-coding genes in *C. briggsae*, those showing a genotype-temperature interaction were significantly enriched in chromosome centers (54.2% of protein-coding interaction genes are on centers compared to 42.9% of those showing no interaction; Fisher’s exact test, *p* < 2.2 x 10^-16^) (Fig. S6).

### Temperature stress weakens the correlation between 22G-RNA and mRNA expression in a genotype-dependent manner

Given the post-transcriptional regulatory role of 22G-RNAs on mRNA for the genes from which they both originate, we tested for changes to the association between 22G-RNA and mRNA expression levels across genotype and temperature treatment combinations. We could directly relate 22G-RNA and mRNA expression for 12,623 genes with data that derived from the same total RNA samples (22G-RNA from this study and mRNA from Mark et al. (2019)). Overall, these 12,623 genes show moderate but significant positive correlations between 22G-RNA and mRNA expression across all genotype-temperature combinations (*p* < 2.2 x 10^-16^ in all genotype-temperature combinations; Fig. S7). The correlation is weakest, however, in temperate HK104 worms reared under both hot and cold chronic thermal stress (Fig. S7; HK104 at 14°C Spearman’s *ρ* = 0.28; HK104 at 30°C Spearman’s *ρ* = 0.23; all other treatment Spearman’s *ρ* ranges from 0.38 to 0.54). Compared to 20°C, the correlation in HK104 is significantly lower at both 14°C (*z*-test, *p* < 2.2 x 10^-16^) and 30°C (*p* < 2.2 x 10^-16^). In AF16 worms, the correlation is also lower at 14°C (*z*-test, *p* < 2.2 x 10^-16^) but higher at 30°C (*p* = 6.7 x 10^-13^), compared to 20°C.

Despite the overall positive correlations, 22G-RNA expression actually correlates negatively with mRNA levels among the 10% of genes with the highest 22G-RNA expression in each genotype-temperature combination (*p* < 6.1 x 10^-16^ in all genotype-temperature combinations; Fig. S7). Again, this pattern is weaker in HK104 at extreme temperatures compared to all other genotype-temperature combinations (Fig. S7; HK104 at 14°C Spearman’s *ρ* = −0.32; HK104 at 30°C Spearman’s *ρ* = −0.22; all other treatment Spearman’s *ρ* ranges from −0.36 to −0.41). Whereas HK104 worms showed a significantly weaker correlation at both 14°C (*z*-test, *p* = 0.021) and 30°C (*p* = 5.9 x 10^-7^) relative to 20°C, AF16 worms did not show significantly different correlations at either 14°C (*p* = 0.260) or 30°C (*p* = 0.602) compared to 20°C. These opposing trends contribute to a nonlinear relationship between 22G-RNA and mRNA expression, showing a peak of mRNA expression for 22G-RNA levels from 5.00 to 6.35 log_2_-RPM (Fig. S8).

Because the 22G-RNA-associated Argonaute protein CSR-1 acts to license and promote mRNA expression whereas the Argonaute WAGO-1 acts to silence mRNA expression (Gu et al. 2009; Conine et al. 2013), we tested for distinct associations of 22G-RNA and mRNA for targets of these Argonautes (see also next section). We found higher mRNA expression of CSR-1 targets than WAGO-1 targets in all genotype-temperature combinations (Fig. S9), consistent with the known roles of CSR-1 and WAGO-1 in promoting and silencing gene expression, respectively, via 22G-RNA targeting (Gu et al. 2009; Conine et al. 2013). As we observed for genes overall, genes targeted by CSR-1 and WAGO-1 both showed a significant positive correlation between 22G-RNA and mRNA expression (Fig. S8; CSR-1 target Spearman’s *ρ* ranges from 0.10 to 0.40 with *p* < 2.5 x 10^-9^ in all genotype-temperature combinations; WAGO-1 target Spearman’s *ρ* ranges from 0.15 to 0.36 with *p* < 2.3 x 10^-5^ in all genotype-temperature combinations). Among genes in the top 10% of 22G-RNA expression for WAGO-1 or CSR-1 targets, WAGO-1 targets tended to have a more negative 22G-RNA: mRNA correlation than CSR-1 targets, consistent with the silencing role of WAGO-1. In particular, at 20°C, the WAGO-1 22G-RNA: mRNA correlation for these genes was significantly negative in HK104 worms but not in AF16 worms (HK104 Spearman’s *ρ* = −0.29, *p* = 0.0076; AF16 *ρ* = −0.15, *p* = 0.160), whereas CSR-1 targets did not show a significant correlation in either HK104 (*ρ* = −0.02, *p* = 0.682) or AF16 (*ρ* = −0.04, *p* = 0.457) animals at this temperature. We observed qualitatively similar but non-significant trends at the more extreme rearing temperatures, suggesting that temperature extremes act to decouple the association between 22G-RNA expression and the expression levels of their targets (WAGO-1 Spearman’s *ρ*: −0.05 to −0.21 with *p* > 0.056; CSR-1 Spearman’s *ρ*: 0.06 to −0.07 with *p* > 0.14).

Upon testing for a shared profile of genotype only, temperature only, or genotype-by-temperature interaction for 22G-RNA and mRNA expression of a given protein coding gene, we detected surprisingly little overlap (Fig. S10). For example, only 30.9% (1269 / 4108) of genes (with both 22G-RNA and mRNA expression data available) showing a 22G-RNA genotype-temperature interaction also showed an mRNA genotype-temperature interaction. This overlap was marginally statistically insignificant (hypergeometric test, *p* = 0.055), and the overlap for other differential expression categories was even lower (Fig. S10). These observations point to the possibility that additional regulatory controls over 22G-RNA and mRNA expression decouples their genotype-by-temperature interactions as reflected in RNA abundances.

### Reduced 22G-RNA expression is specific to the EGO-1/CSR-1 pathway

RNA-dependent RNA polymerases (RdRPs), such as EGO-1 and RRF-1, initiate the biogenesis of 22G-RNAs that then bind to Argonaute proteins, including CSR-1, WAGO-1, and HRDE-1. These Argonautes serve as effectors that modulate expression of target genes in the germline, mediated by base complementary binding with 22G-RNA sequences. CSR-1 binds 22G-RNAs made by EGO-1, WAGO-1 binds 22G-RNAs made predominantly by RRF-1 and by EGO-1 to a lesser extent, and HRDE-1 binds 22G-RNAs made by both EGO-1 and RRF-1 (Claycomb et al. 2009; Gu et al. 2009; Buckley et al. 2012) (Fig. 5B). To investigate these 22G-RNA pathway proteins as potential sources of temperature-dependent reductions in 22G-RNA expression, we quantified their mRNA expression profiles. *Cbr-ego-1*, *Cbr-rrf-1*, *Cbr-csr-1*, *Cbr-wago-1*, and *Cbr-hrde-1* all shared similar mRNA expression across all six genotype-temperature combinations (Fig. 5A). In the tropical AF16 strain, mRNA expression of these five genes was insensitive to rearing temperature and, at 20°C, mRNA expression was indistinguishable between AF16 and HK104. The stressful rearing temperatures of both 14°C and 30°C, by contrast, typically led to reduced expression of these RdRPs and Argonautes in the temperate HK104 strain (Fig. 5A). We also profiled the mRNA expression of several genes in the 26G-RNA and piRNA pathways, which act upstream of 22G-RNA production (Hoogstrate et al. 2014), although these did not generally show a consistent pattern (Fig. S11)

**Figure 5.**
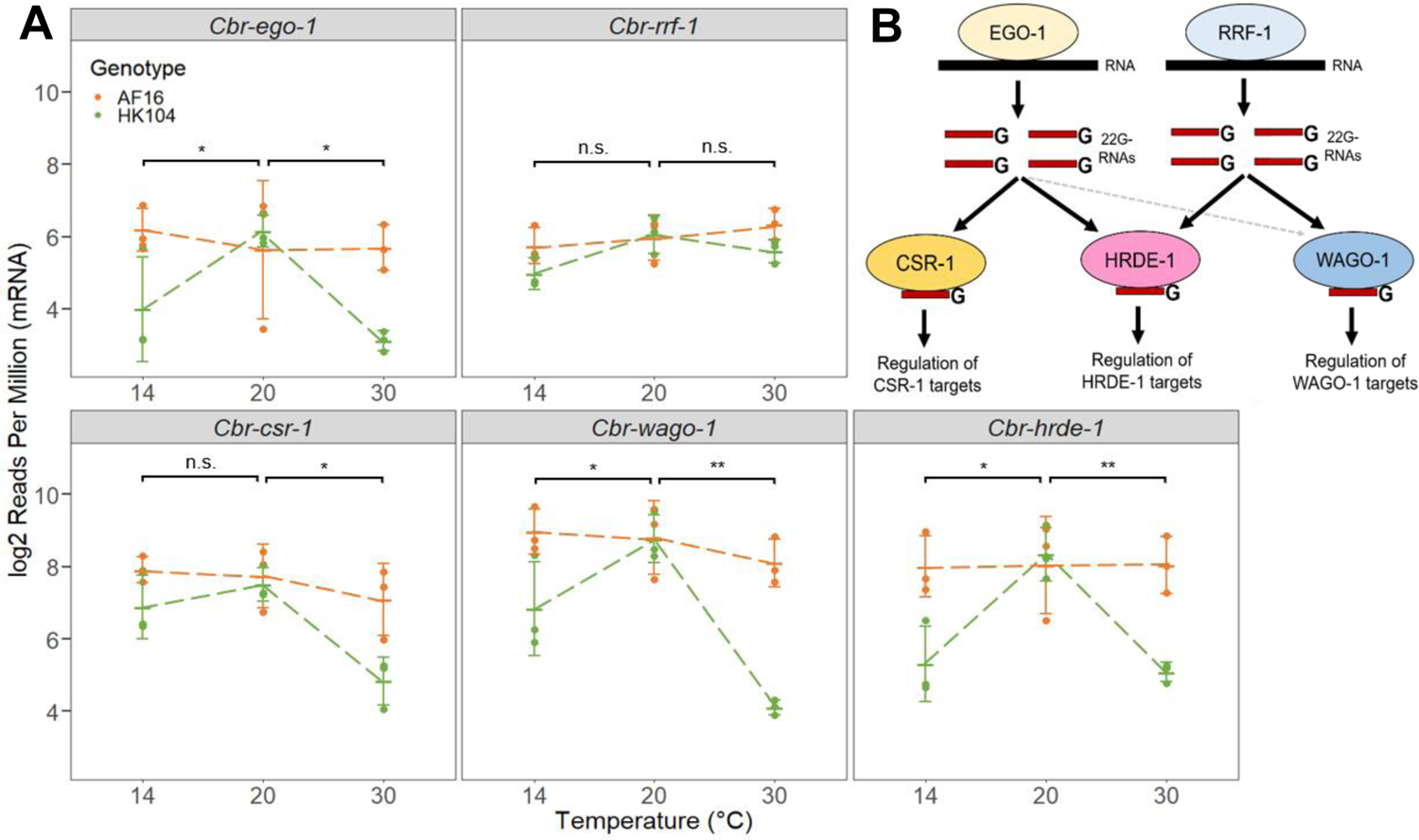
(A) mRNA expression of different genes involved in the 22G-RNA pathway, across genotypes and temperatures. Error bars give ±1 standard deviation around the mean for each set of replicates. Dashed lines connect mean expression values for each genotype across temperatures. A linear model was fit to determine the significance of genotype-temperature interactions between 14°C and 20°C, and between 20°C and 30°C, with * indicating *p* < 0.05, ** indicating *p* < 0.01, and n.s. indicating non-significance. (B) Summary of the 22G-RNA pathway proteins studied here. The RNA-dependent RNA polymerases (RdRPs) EGO-1 and RRF-1 produce 22G-RNAs, which are bound by the Argonaute proteins CSR-1, WAGO-1, and HRDE-1. CSR-1-bound 22G-RNAs are generally produced by EGO-1, whereas WAGO-1-bound 22G-RNAs are mainly produced by RRF-1 but also EGO-1 to some degree. Both EGO-1 and RRF-1 produce 22G-RNAs bound by HRDE-1.

These observations point to the possibility that the differences in 22G-RNA expression (Fig. 3, Fig. 4) arise from changes in mRNA expression of genes in the 22G-RNA pathway. Reductions to 22G-RNA expression should be strongest in the genes targeted by a given pathway if changes to the pathway’s Argonaute or RdRP expression cause lower 22G-RNA expression. To explore this possibility, we first compiled sets of genes targeted by the 22G-RNAs associated with EGO-1, RRF-1, CSR-1, WAGO-1, or HRDE-1 from previous RNA-sequencing studies in *C. briggsae* (for CSR-1) or *C. briggsae* orthologs of *C. elegans* targets (for EGO-1, RRF-1, WAGO-1, HRDE-1, and the CSR-1a and CSR-1b isoforms of CSR-1) (Claycomb et al. 2009; Vasale et al. 2010; Buckley et al. 2012; Tu et al. 2015; Charlesworth et al. 2021). The resulting putative target sets contain between 146 and 4839 genes (median 1706 genes), for which we obtained *C. briggsae* 22G-RNA expression data of 121 to 3953 genes (median 1456 genes) (Table 2; Table S4). Consistent with their function in shared 22G-RNA pathways, EGO-1 and CSR-1 show extensive target overlap (1427 out of 1706 EGO-1 targets are also CSR-1 targets, 83.6%), as do RRF-1 and WAGO-1 (72 out of 146 RRF-1 targets are also WAGO-1 targets, 49.3%) (Fig. S12). We observed that 76.8% (726 / 945) of HRDE-1 targets were also either CSR-1 targets (368 / 945) or WAGO-1 targets (395 / 945), affirming HRDE-1’s participation in both the EGO-1/CSR-1 and RRF-1/WAGO-1 pathways (Fig. S12).

**Table 2:**
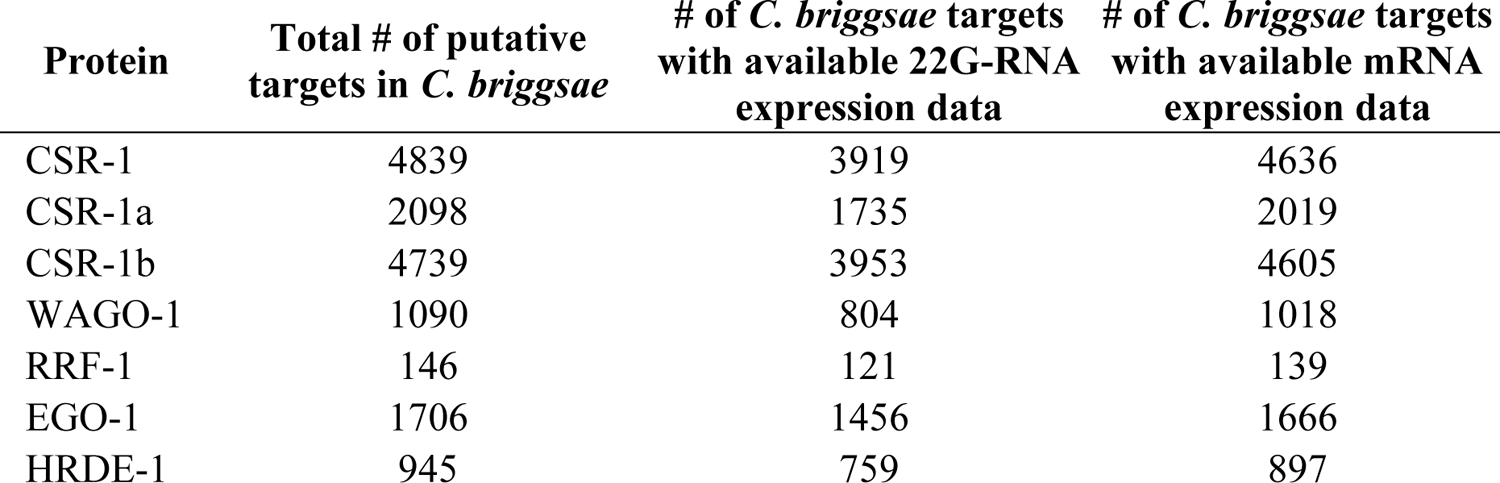
Number of 22G-RNA pathway protein targets in *C. briggsae* used in this study. CSR-1 targets are known for *C. briggsae*; targets of all other proteins are orthologs of known targets in *C. elegans*.

| **Protein** | **Total # of putative targets in *C. briggsae*** | **# of *C. briggsae* targets with available 22G-RNA expression data** | **# of *C. briggsae* targets with available mRNA expression data** |
| --- | --- | --- | --- |
| CSR-1 | 4839 | 3919 | 4636 |
| CSR-1a | 2098 | 1735 | 2019 |
| CSR-1b | 4739 | 3953 | 4605 |
| WAGO-1 | 1090 | 804 | 1018 |
| RRF-1 | 146 | 121 | 139 |
| EGO-1 | 1706 | 1456 | 1666 |
| HRDE-1 | 945 | 759 | 897 |

Comparing the 22G-RNA expression of the targets of EGO-1/CSR-1, HRDE-1, and RRF-1/WAGO-1 between genotypes and temperatures revealed striking differences between the pathways. Genes targeted by EGO-1 and CSR-1 showed 22G-RNA profiles that mirrored the genome-wide pattern: compared to 20°C, 22G-RNA expression was significantly reduced specifically at 30°C in HK104 but not AF16 worms (Fig. 6), with AF16 worms showing slightly increased expression at 30°C. While both CSR-1a and CSR-1b targets shared this same trend, it was most pronounced for CSR-1b targets, and genes exclusively targeted by CSR-1a and not CSR-1b did not show this trend at all (Fig. S13). These observations implicate the germline-constitutive CSR-1b short isoform, and not the intestinal and spermatogenesis-associated CSR-1a long isoform (Charlesworth et al. 2021), in this pattern of genotype-dependent differential expression of 22G-RNAs at 30°C.

**Figure 6.**
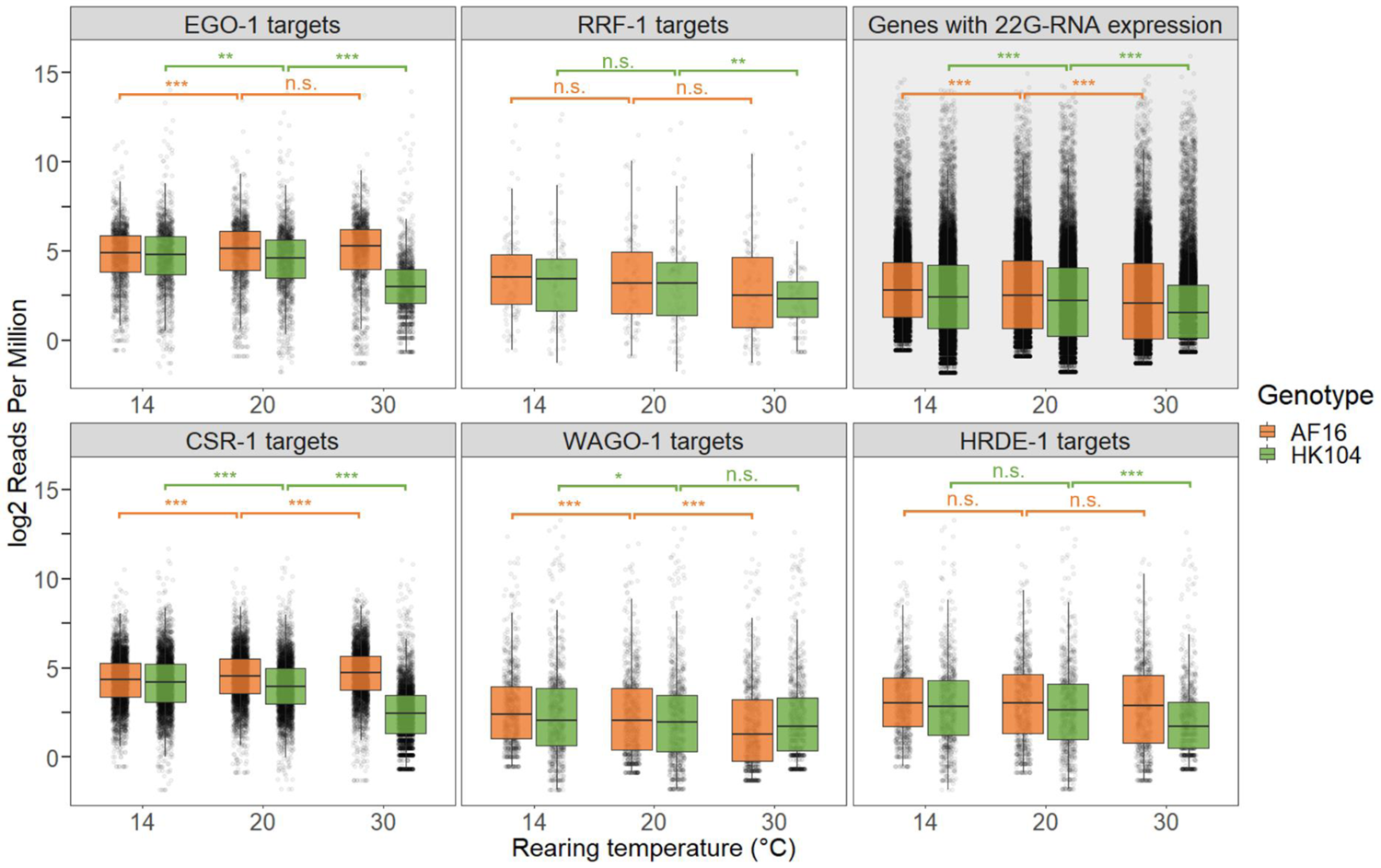
22G-RNA expression of genes targeted by EGO-1, RRF-1, CSR-1, WAGO-1, or HRDE-1, as well as all genes showing 22G-RNA expression, for each genotype-temperature combination. Significant differences between temperature treatments of the same genotype were determined using a Wilcoxon rank sum test, with * indicating *p* < 0.05, ** indicating *p* < 0.01, *** indicating *p* < 0.001, and n.s. indicating non-significance.

Genes targeted by HRDE-1 showed the same overall pattern as EGO-1 and CSR-1 targets, but weaker in magnitude, with high temperatures leading to significantly reduced 22G-RNA expression in HK104, but not AF16 (Fig. 6). This pattern is likely due to partial overlap between EGO-1 and HRDE-1 targeting, as 37.5% (354 / 945) of HRDE-1 targets are also EGO-1 targets. Indeed, partitioning HRDE-1 targets into those genes also targeted by EGO-1 and those not targeted by EGO-1 showed that the decrease in HK104 worms at 30°C was strongest in HRDE-1 targets that are also EGO-1 targets (Fig. S14). Surprisingly, genes targeted by RRF-1 and WAGO-1 did not show the genome-wide pattern. Instead, 22G-RNA expression was either the same (RRF-1 targets) or higher (WAGO-1 targets) at 30°C in HK104 compared to AF16 (Fig. 6), though at 30°C RRF-1 targets showed a significant decrease in HK104 but not AF16, similar to HRDE-1 targets. These profiles of target gene 22G-RNA expression match the profiles of target gene mRNA expression across genotypes and temperatures, including a broad decrease in mRNA expression for EGO-1 and CSR-1 targets at 30°C in HK104 (Fig. S15), consistent with the overall positive association between 22G-RNA and mRNA expression levels.

Our gene-level counts of 22G-RNA expression were normalized without using the standard trimmed mean of M-values (TMM) method to avoid the assumption that most genes do not show differential expression (Evans et al. 2018), as our data violated this assumption. Nevertheless, these results are robust to TMM normalization. Applying TMM scaling to the 22G-RNA expression of these 14,283 genes, we find that targets of EGO-1 and CSR-1 (but not HRDE-1) still show greatly decreased 22G-RNA expression in HK104 but not AF16 worms at 30°C despite the normalization removing this pattern from the background distribution of genes (Fig. S16). Together, these results support the idea that temperature stress at 30°C uniquely disrupts the germline-expressed EGO-1 RdRP/CSR-1 Argonaute pathway in the temperate HK104 genotype. This occurs primarily through the CSR-1b isoform, and depresses the expression of hermaphrodite germline-associated 22G-RNAs and their target genes.

## Discussion

Strains of *C. briggsae* from tropical and temperate latitudes show responses to temperature stress that are consistent with local adaptation: tropical strains are more fecund than temperate strains when raised at high temperatures, whereas temperate strains are more fecund and viable than tropical strains when exposed to low temperatures (Prasad et al. 2011; Wang et al. 2021). Their responses are also genetically distinct in terms of behavior (Stegeman at al. 2013; Stegeman et al. 2019), mRNA expression (Mark et al. 2019; Wang et al. 2021), and in sperm fertility and germline development (Prasad et al. 2011; Poullet et al. 2015). Here, we incorporate differentiation in the expression of endogenous small RNAs by sequencing small RNAs from tropical and temperate *C. briggsae* genotypes raised under distinct temperature conditions. We find that some temperature-induced changes in small RNA expression are common to both genotypes, such as reduced piRNA expression, whereas other changes are genotype-specific. In particular, we document how globally reduced 22G-RNA expression is specific to the temperate HK104 genotype at a high 30°C rearing temperature, possibly helping to explain why high temperatures have deleterious fitness consequences on this genetic background. Thus, we find that the evolution of small RNA pathways may be a contributor to the local adaptations to temperature seen in *C. briggsae*. While much is unknown about the role of small RNAs (and endo-siRNAs in particular) in local adaptation, our work suggests that these pathways are potentially important and warrant further study in this and other species.

Both the tropical AF16 genotype and the temperate HK104 genotype showed decreased overall piRNA expression at high temperatures, in line with similar findings in *C. elegans* and other *C. briggsae* strains (Belicard et al. 2018). The three distinct genomic clusters of piRNA loci (Shi et al. 2013), however, showed differing expression responses to extreme temperatures. The piRNAs encoded in the cluster on the left arm of chromosome IV (0.6 Mbp – 7.0 Mbp) showed decreased expression at 30°C, whereas those on the right arm (13.2 Mbp – 15.0 Mbp) showed increased expression at 14°C. It is currently not known whether these clusters participate in different silencing pathways or target different sets of transcripts in *C. briggsae*, and so further work is needed to determine the full significance of this cluster-dependent piRNA expression. In nematodes, piRNAs act in the germline to silence transposons and other foreign DNA sequences (Lee et al. 2012), and disruptions of this pathway cause fertility defects at high temperatures (Wang and Reinke 2008). However, the role of piRNAs in cold temperature phenotypic responses has yet to be characterized. The cluster-specific expression changes of *C. briggsae* piRNAs at extreme temperatures in both genotypes may therefore point to a role associated with the general reduction in lifetime fecundity seen at these temperatures (Prasad et al. 2011).

For 22G endo-siRNAs, the most common type of small RNA in our samples, we observed the largest expression changes for the temperate HK104 strain when reared at 30°C. We documented substantial genome-wide reductions in 22G-RNA expression in this genotype-by-temperature combination for multiple different types of loci, on multiple different chromosomes, and at numerous different loci across the full length of each chromosome. Together, these observations point to a global, genome-wide decrease in expression that is specific to the HK104 genotype when reared under chronic high thermal stress. These broad changes in 22G-RNA expression may contribute to the significant decrease in lifetime fecundity seen in the temperate HK104 strain when reared at 30°C (Prasad et al. 2011); disruptions to *C. briggsae* 22G-RNA pathways are known to have a negative effect on offspring viability (Tu et al. 2015). Evidence from mRNA sequencing suggests that many genes are indeed misregulated in the HK104 strain under heat stress (Mark et al. 2019), although it remains unclear to what extent perturbed mRNA expression levels depend on transcriptional versus post-transcriptional regulatory mechanisms, such as 22G-RNA-associated Argonaute pathways.

On average, genes targeted by the Argonaute CSR-1 and its RNA-dependent RNA polymerase EGO-1 showed reduced expression of antisense 22G-RNAs at high temperatures in the temperate HK104 strain but not the tropical AF16 strain. We generally did not observe this genotype-temperature interaction, however, for targets of the RRF-1/WAGO-1 pathway. This difference in 22G-RNA target profiles may arise from differences in gene expression of the RdRPs: *Cbr-ego-1* showed substantially reduced mRNA expression at high temperatures in only HK104, whereas *Cbr-rrf-1* showed a weaker reduction. As EGO-1 is needed for the biogenesis of the 22G-RNAs bound by CSR-1 (Claycomb et al. 2009), downregulation of EGO-1 expression in this genotype under heat stress likely leads to the production of fewer 22G-RNAs that can bind to CSR-1. We also observe decreased mRNA expression for EGO-1/CSR-1 targets in HK104 at high temperatures, as expected given that CSR-1 plays a positive role in promoting gene expression of its targets (Conine et al. 2013; Wedeles et al. 2013).

The overall positive association between 22G-RNA expression and mRNA expression, however, makes it difficult to conclude whether decreased 22G-RNA expression causes decreased mRNA expression or vice versa. For instance, at 30°C, targets of WAGO-1 show increases in both 22G-RNA expression and mRNA expression in HK104 compared to AF16, despite WAGO-1’s role in gene silencing (Gu et al. 2009). Nevertheless, large-scale misregulation of EGO-1/CSR-1 targets in the germline may partially explain the reduced sperm fertility that temperate *C. briggsae* strains show at elevated temperatures (Prasad et al. 2011). If so, then temperate *C. briggsae* strains may have acquired mutations involving the EGO-1/CSR-1 pathway that compromise the adaptive high temperature tolerance of tropical strains and ancestral genotypes – perhaps by causing the downregulation of EGO-1 at high temperatures – although further study is needed to test possible mechanisms directly.

*C. briggsae* and other *Caenorhabditis* nematodes express both a long isoform (CSR-1a) and a short isoform (CSR-1b) of CSR-1, which differ due to an additional 163 N-terminal amino acids in CSR-1a (Tu et al. 2015; Charlesworth et al. 2021). We found that HK104 worms showed reduced 22G-RNA expression at 30°C compared to 20°C in targets of both CSR-1 isoforms, though this trend was more pronounced in CSR-1b targets than CSR-1a targets. In *C. elegans*, genes targeted by only the CSR-1b isoform are enriched for EGO-1 targets whereas genes targeted by only the CSR-1a isoform are enriched for RRF-1 targets (Charlesworth et al. 2021; Nguyen and Philips 2021). Consistent with our conclusion that EGO-1 and not RRF-1 is the key RdRP associated with the genotype-specific 22G-RNA reduction at 30°C, we did not observe this 22G-RNA reduction in the genes targeted only by CSR-1a. Interestingly, expression of CSR-1a is required for full fertility under high temperature stress in *C. elegans* due to its role in spermatogenesis (Charlesworth et al. 2021). As CSR-1a is only expressed in the germline during the L4 larval stage when spermatogenesis occurs (Charlesworth et al. 2021), it is possible that our RNA sequencing of adult worms has missed temperature-dependent differences between strains in the CSR-1a pathway associated with spermatogenesis.

One limitation to our analyses of the different 22G-RNA pathways is that the sets of EGO-1 and RRF-1 targets used here are almost certainly incomplete. For instance, 70.0% (3388 / 4839) of the genes we defined as CSR-1 targets were not also defined as either EGO-1 or RRF-1 targets, despite the fact that EGO-1 is thought to be the main source of 22G-RNAs bound by CSR-1 (Claycomb et al. 2009). Part of this incompleteness is simply a byproduct of the experimental approaches used to define them, as the single-gene mutants used may not be sufficient in recovering the full set of EGO-1 or RRF-1 target genes, given that EGO-1 and RRF-1 are partially redundant in 22G-RNA biogenesis (Claycomb et al. 2009; Gu et al. 2009; Vasale et al. 2010). In addition, we relied on orthology of gene targets between *C. elegans* and *C. briggsae* for most Argonaute pathway members. Comprehensive profiling of EGO-1 and RRF-1 targets in *C. briggsae* itself will benefit future analysis for how these RdRPs may influence *C. briggsae* local adaptation.

We found that 22G-RNA expression correlated positively with mRNA expression in all genotype-temperature combinations, even among genes targeted by WAGO-1, which is thought to silence its targets (Gu et al. 2009). This general positive correlation is likely due to the requirement of mRNAs for the biogenesis of complementary 22G-RNAs, as these mRNAs serve as a template utilized by RdRPs in 22G-RNA production (Hoogstrate et al. 2014). However, we found that genes with the highest levels of 22G-RNA expression, including WAGO-1 targets, negatively correlated with mRNA expression, consistent with 22G-RNA-mediated silencing at sufficient levels of 22G-RNA expression. This non-linear relationship between 22G-RNA and mRNA expression also has been observed in the *C. elegans* germline (Bezler et al. 2019).

Surprisingly, however, we observed little overlap between genes with genotype- or temperature-dependence in 22G-RNA expression and those genes showing the same effects for mRNA expression. This disparity could suggest that most changes in 22G-RNA expression are not strong enough to perturb mRNA expression in a way that matches the 22G-RNA profile, especially if pre-transcriptional regulation (e.g., chromatin modifications that regulate transcription initiation) is a potent factor in controlling genotype- or temperature-dependent changes in transcription. Another potential explanation is that 22G-RNAs may disproportionately regulate target gene expression through processes downstream of mRNA. For example, endo-siRNAs in plants that are 22 nucleotides in length are capable of repressing the mRNA translation of target genes (Wu et al. 2020). Similar translation inhibition may also occur in some genes showing 22G-RNA effects in *C. briggsae*, although data supporting a link between 22G-RNAs and translational regulation in *Caenorhabditis* are lacking.

Overall, we find that tropical and temperate strains of *C. briggsae* show significant differences in how endogenous small RNA pathways respond to the same chronic temperature stress. Heat stress induced a genome-wide decrease in the expression of endogenous 22G-RNAs, but only in the strain derived from a temperate latitude. Genotype-specific differences in 22G-RNA expression, potentially caused by altered expression of genes involved in CSR-1b-associated 22G-RNA biogenesis such as EGO-1, may contribute to heightened sensitivity to heat stress with profound consequences for animal fitness. *C. briggsae* thus provides a powerful experimental system to elucidate how small RNA mechanisms may contribute to local adaptations in nature.

## Materials and Methods

### Small RNA sequencing datasets

To assess the sensitivity of small RNA expression to rearing temperature and genetic background, we isolated and sequenced small RNAs for two isogenic strains of *C. briggsae*, each reared under defined temperature conditions throughout development (14°C, 20°C, and 30°C). The AF16 strain represents a tropical latitude genotype whereas HK104 represents a genetically differentiated temperate latitude genotype, with the extreme 14°C and 30°C temperatures representing chronic stressful conditions for both genotypes (Cutter et al. 2006; Prasad et al. 2011; Félix et al. 2013; Thomas et al. 2015). Worms were cultured as described in Mark et al. (2019). Briefly, worms were plated on 100mm Nematode Growth Medium plates with 1.5mL of concentrated OP50 *Escherichia coli*. Plates were incubated at 14°C, 20°C, and 30°C for approximately 150 hrs, 65 hrs and 48 hrs, respectively, to the mature adult stage. Total RNA from gravid adult hermaphrodites in mass isogenic cultures of each strain-temperature combination was isolated with TRIzol extraction and isopropanol precipitation (Tu et al. 2015) from three biological replicates, for a total of 18 distinct samples that correspond to the identical samples analyzed for mRNA expression by Mark et al. (2019). Using the Ambion MirVana kit, we separated the small RNA fractions <200 bp and then isolated small RNAs between 18 and 30 nucleotides using gel extraction from 15% acrylamide gels, as adapted from Claycomb et al. (2009) and Gu et al. (2011). The samples were treated with tobacco acid pyrophosphatase to convert triphosphorylated RNA into a monophosphorylated form prior to ligating the 3’ linker, followed by 15% polyacrylamide/7M urea gel extraction (Claycomb et al. 2009; Gu et al. 2011). A 5’ linker sequence, possessing one of nine 4-nucleotide barcodes at its 3’ end, was ligated to the samples to enable multiplexing. This was followed by reverse-transcription cDNA synthesis, and PCR amplification, to add Illumina sequencing adapters with an additional 8-nucleotide indexing barcode on the 3’ adapter.

Small RNA libraries from each replicate were sequenced at the Tufts University Core Facility. Samples containing 202,528,982 total raw reads from AF16 and HK104 were demultiplexed and had their 5’ 4-nucleotide barcodes removed using a custom Perl script. Before barcode removal, Cutadapt v1.2.1 (Martin 2011) was used to remove 3’ adapters (parameters: -e0.1 -m 21 -M 34 -a CTGTAGGCACCATCAAT) and discard reads that contained at least 14 nucleotides of the 5’ adapter (parameters: -e 0.1 --untrimmed-output= -O 14 -g TCTACAGTCCGACGATC). Reads were discarded if, after adapter and barcode trimming, the read was longer than 30 nucleotides or shorter than 17 nucleotides. After trimming, the 18 samples averaged 9,660,349 total reads each, giving an average of 28,981,046 reads for each of the six treatments (Table S5). Analysis of all samples using FastQC v0.11.8 (Andrews 2010) confirmed for 17 of them that FASTQ files contained, as expected, a majority of reads of approximately 22 bases in length and no detected adapter sequences. However, one replicate of HK104 grown at 30°C showed an unusually low read count (1,298,689 vs >4,500,000 reads in all other samples; Table S5) with a length distribution skewed heavily towards short 17-19 nucleotide reads. These anomalies led us to exclude this sample from downstream analyses other than genome alignment (see below), resulting in an average of 10,152,211 reads per sample and 28,764,597 reads per treatment.

### Generation of a genotype-specific reference for HK104 worms

To align these small RNAs to the *C. briggsae* genome for expression quantification, we used the reference genome sequence and gene annotations from WormBase release WS272 (Lee et al. 2018). Since this reference genome is based on the tropical AF16 strain, we also generated a genotype-specific reference for the temperate HK104 strain. Whole-genome sequencing reads for the HK104 strain from Mark et al. (2019) were obtained from NCBI’s Sequence Read Archive (accession SRR8333803) and aligned to the AF16 reference genome. Alignments were performed with NextGenMap v0.5.2 (Sedlazeck et al. 2013) in paired-end mode, with all other parameters left as default, including the default cutoff of 65% alignment identity to consider a read as successfully mapped. NextGenMap is able to account for a high level of divergence between the sequences being aligned, making it suitable for performing alignments between different strains. These alignments were used to call single nucleotide polymorphisms (SNPs) and short indels between the AF16 and HK104 genotypes with the HaplotypeCaller function from the GATK v4.1.4.1 software toolkit run with default parameters (McKenna et al. 2010). Using a custom Python script, we substituted these SNPs and indels into the AF16 reference genome sequence in order to create an HK104 “pseudo-reference” genome sequence. We then ran flo (Pracana et al. 2017) with default parameters to create a gene annotation file for HK104 by transferring gene annotations (microRNA genes, protein-coding genes, pseudogenes, transposable elements, repeat regions, rRNA genes, tRNA genes, and other noncoding RNA genes) from the WormBase AF16 reference to the coordinates of the HK104 pseudo-reference sequence (available at https://github.com/Cutterlab/C_briggsae_small_RNAs). HK104 protein-coding gene annotations were kept only for 17,038 genes (81.8% of 20,829 protein-coding genes total) in which all constituent parts (e.g. exons, UTRs) could be successfully converted from AF16 coordinates to HK104 coordinates.

### Alignment of reads to reference genomes

Small RNAs were aligned to the *C. briggsae* genome using the STAR v2.7.2a read alignment software (Dobin et al. 2013). All small RNA samples were aligned to both the AF16 genome (using the genome sequence and gene annotations from WormBase) and the HK104 genome (using the genotype-specific genome sequence and gene annotations created here). Both genomes were compiled with STAR’s genomeGenerate function using the following options: --genomeSAindexNbases 12 --sjdbOverhang 29. Small RNA alignments to each genome were run with the following options: --quantMode GeneCounts --limitBAMsortRAM 30000000000 --outSAMtype BAM SortedByCoordinate --outFilterMultimapNmax 50 --outFilterMultimapScoreRange 0 --outFilterMismatchNoverLmax 0.05 --outFilterMatchNmin 16 --outFilterScoreMinOverLread 0 --outFilterMatchNminOverLread 0 --alignIntronMax 1. AF16 small RNA reads mapped best to the AF16 reference genome and HK104 reads mapped best to our HK104 pseudo-reference genome (Table S6). Therefore, we only used alignments of small RNAs to their corresponding genome for further analyses. The number of reads from each replicate that aligned to their corresponding genome are given in Table S5.

### Counting aligned small RNA reads

Aligned small RNA reads from all samples were counted using a custom R pipeline (Charlesworth et al. 2021; https://github.com/ClaycombLab/Charlesworth_2020). Mapped reads were counted only if they aligned to the genome (either sense or antisense) without any insertions, deletions, or mismatches, but allowing for gaps due to introns. Each perfectly-aligned read was annotated with its length, first nucleotide, and coordinate location in the genome, using the location of the primary alignment (as determined by STAR) for reads that aligned to multiple locations. For each annotated genomic feature on *C. briggsae*’s six chromosomes (microRNA genes, protein-coding genes, pseudogenes, transposable elements, repeat regions, rRNA genes, tRNA genes, other noncoding RNA genes, and a single feature comprising all unannotated regions), we counted how many perfectly-aligned reads of each length and first nucleotide aligned completely to this feature in each replicate, separately for sense alignments and antisense alignments (Table S5). For reads that aligned perfectly to multiple different locations in the genome at the same time, each alignment of that read to a feature was only counted as 1/N reads for that feature, where N is the total number of times the read aligned to a feature of that class (e.g. protein-coding genes).

These counts were used to determine the total number of microRNAs, 21U-piRNAs, and 22G and 26G endo-siRNAs in each replicate expressed from each *C. briggsae* chromosome, including unplaced genomic scaffolds. We defined microRNAs as reads aligning to the sense strand of annotated microRNA genes, 21U-piRNAs as any reads with a length of exactly 21 bases beginning with a uridine (U) nucleotide, and 22G- and 26G-RNAs as any reads with a length between 21-23 bases and 25-27 bases, respectively, and beginning with a guanine (G) nucleotide. Due to their observed numerical abundance, we primarily focused on characterizing the expression of 22G-RNAs. The total number of 22G-RNAs aligning to each of the six *C. briggsae* chromosomes (excluding unplaced genomic scaffolds) was fit using the linear model: 22G-RNA expression = chromosome of origin effect + genotype effect + temperature effect + genotype:temperature interaction effect. This linear model was used to assess the significance of the temperature effect for each genotype and the genotype-temperature interaction, at 30°C relative to 20°C, as well as the chromosome effect for each chromosome. All statistical analyses were done in R v4.1.1 (R Core Team 2021). Coordinates for chromosome arm and center domains in the current *C. briggsae* genome assembly (cb4) were taken from Ross et al. (2011), combining the short chromosome tip regions with the adjacent arm domains.

### 22G-RNA expression analysis

To investigate the expression of 22G endo-siRNAs across genotypes and temperature treatments, we quantified the number of 22G-RNAs aligning antisense to all annotated repeats, transposons, pseudogenes, and protein-coding genes in the *C. briggsae* genome, using only the 16,916 unique features whose coordinates completely transferred from the AF16 genome annotation to the HK104 genome annotation. Repeat and transposon classes were kept if at least one copy of the repeat/transposon was present in both genome annotations (i.e. without requiring every copy to be present in both genomes). The total number of 22G-RNAs aligning antisense to these 16,916 features was fit, separately for protein-coding genes, pseudogenes, repeats, and transposons, using the linear model: 22G-RNA expression = genotype effect + temperature effect + genotype:temperature interaction effect. These linear models were used to assess the significance of the temperature effect for each genotype and the genotype-temperature interaction, at 30°C relative to 20°C. These 16,916 features were filtered to exclude those without at least 1 Read Per Million (RPM) of 22G-RNA expression in at least 3 different replicates, resulting in a final set of 14,283 features (13,668 protein-coding genes, 388 pseudogenes, 187 repeats, and 40 transposons) used for further analysis. To assess the similarity of 22G-RNA expression across the 17 samples, we created a multidimensional scaling plot of replicates based on the 22G-RNA expression of these 14,283 genomic features using the plotMDS function from the limma R package (Ritchie et al. 2015). 22G-RNA expression values were normalized to units of log_2_ RPM using voom (Law et al. 2014), implemented in the limma R package (Ritchie et al. 2015). We did not include trimmed mean of M-values (TMM) normalization in our normalization pipeline, as our 22G-RNA expression data violate the assumption of TMM that most genes are not differentially expressed, which can lead to inflated false positive rates (Evans et al. 2018).

Features with differential 22G-RNA expression between genotype-temperature combinations were detected using limma with the following linear model: 22G-RNA expression = genotype effect + temperature effect + genotype:temperature interaction effect. Features with an adjusted *p*-value below a false-discovery rate of 0.05 were considered significant. We then classified features into five non-overlapping groups based on the presence of significant genotype, temperature, or interaction effects: 1) “genotype only” features have a significant genotype effect, but no significant temperature effect or interaction, 2) “temperature only” features have a significant temperature effect, but no significant genotype effect or interaction, 3) “additive genotype-temperature” features have significant genotype and temperature effects, but no significant interaction, 4) “genotype-temperature interaction” features have a significant interaction effect, and 5) “no effect” features do not have a significant genotype, temperature, or interaction effect. For each of these five sets of features, we plotted voom-normalized small RNA expression across all 17 samples using the ComplexHeatmap R package (Gu et al. 2016). Expression values for each heatmap row were row-normalized using R’s scale function (R Core Team 2021) and ordered by hierarchical clustering. The distribution of protein-coding genotype-temperature interaction genes across each chromosome was visualized using the density calculated by R’s hist function (R Core Team 2021), using a bin size of 250kb.

### Comparisons with mRNA expression

We processed mRNA expression data from Supplementary File S1 of Mark et al. (2019) that correspond to each of the 17 *C. briggsae* replicate samples used here. This dataset contains voom-normalized mRNA expression data for 16,199 different protein-coding genes, along with classifications describing whether each gene shows significant genotype, temperature, or interaction effects on mRNA expression. We compared expression levels for the 12,623 genes with both 22G-RNA and mRNA expression values available for each replicate in all six genotype-temperature combinations (regardless of expression level), using the mean expression of all replicates in a given treatment. We did not further normalize these gene-level values of 22G-RNA and mRNA expression to log_2_ Reads Per Kilobase Million (RPKM), as doing so resulted in significant negative correlations between 22G-RNA/mRNA expression and gene length, although our conclusions with respect to mRNA and 22G-RNA expression do not qualitatively change with normalization to log_2_ RPKM rather than log_2_ RPM (data not shown). For scatterplots of 22G-RNA versus mRNA expression for these genes, we fit a second-order polynomial curve to each plot and first calculated the correlation between 22G-RNA and mRNA expression of all 12,623 of these genes using Spearman’s rank correlation coefficient. We then computed the correlation again using only the genes in the top 10% of 22G-RNA expression in each genotype-temperature combination. To test whether these correlation coefficients were significantly different from 0, we used the cor.test function in R (R Core Team 2021). The significance of differences between correlation coefficients was calculated using a *z*-test after Fisher’s *z*-transformation (Myers and Sirois 2006). The significance of the overlap between the genes showing a genotype-temperature interaction for both 22G-RNA and mRNA expression was calculated using a hypergeometric test and visualized using the VennDiagram R package (Chen and Boutros 2011).

### Expression of small RNA pathway proteins and targets

To associate trends in 22G-RNA expression with the mRNA expression of proteins involved the 22G-RNA pathway, we extracted Argonaute and RNA-dependent RNA polymerase (RdRP) expression profiles for *C. briggsae* from the Mark et al. (2019) dataset. To assess the significance of the genotype-temperature interactions on mRNA expression for these genes between 14°C and 20°C, and between 20°C and 30°C, we modeled voom-normalized mRNA expression for each gene using the linear model: 22G-RNA expression = genotype effect + temperature effect + genotype:temperature interaction effect.

Next, we compiled the identity of genes targeted by these proteins from prior studies of small RNA sequencing that used either immunoprecipitation or mutation of the protein of interest. CSR-1 targets, as determined by CSR-1 immunoprecipitation in *C. briggsae*, were taken from Supplementary Table S2 of Tu et al. (2015). For other 22G-RNA pathway proteins (CSR-1 isoforms CSR-1a and CSR-1b, WAGO-1, RRF-1, EGO-1, and HRDE-1), target genes were defined as the *C. briggsae* orthologs of target genes from experiments conducted in *C. elegans*, using the *C. briggsae* – *C. elegans* orthologs defined in Tu et al. (2015). Given that 79.6% (3851 / 4839) of CSR-1 targets in *C. briggsae* are orthologs of CSR-1 targets in *C. elegans* (Tu et al. 2015), we expect this orthology approach to be valid presuming that other small RNA pathways show similarly high levels of target conservation between species. This orthology-based approach included genes targeted in *C. elegans* by the CSR-1 isoforms CSR-1a and CSR-1b (taken from “IP Enrichment (22G reads)” in Supplementary Table S2 from Charlesworth et al. (2021)), WAGO-1 (Supplementary Table S2 of Tu et al. (2015)), RRF-1 (“*rrf-1*” in Supplementary Table S4 of Vasale et al. (2010)), EGO-1 (Supplementary Table S5 of Claycomb et al. (2009)), and HRDE-1 (genes in “GeneList_FINN1top500.txt” and “Sheet2” of Supplementary Table 2 from Buckley et al. (2012) with an RPKM of at least 50). Wilcoxon rank sum tests were used to calculate the significance of differences in voom-normalized 22G-RNA and mRNA expression of target genes between temperature treatments of the same genotype.

## Acknowledgements

We thank members of the Cutter lab for providing thoughtful feedback on an earlier version of this manuscript. We also thank Wei Wang, Santiago Sánchez-Ramírez, and Andrew Lugowski for providing scripts used in some analyses. This work was supported by the Natural Sciences and Engineering Research Council of Canada to A.D.C. (RGPIN-2018-05098) and to J.M.C (RGPIN-418683 and RGPIN-2020-06235) and the Canadian Institutes of Health Research to J.M.C. (Canada Research Chair Tier II in Small RNA Biology).

## Data availability

Small RNA sequencing data is available in Gene Expression Omnibus (GEO), under accession number GSE202589. Scripts used to perform the analyses described here, as well as the pseudo-reference genome sequence for HK104, are available at https://github.com/Cutterlab/C_briggsae_small_RNAs.

---

**Figure S1.**
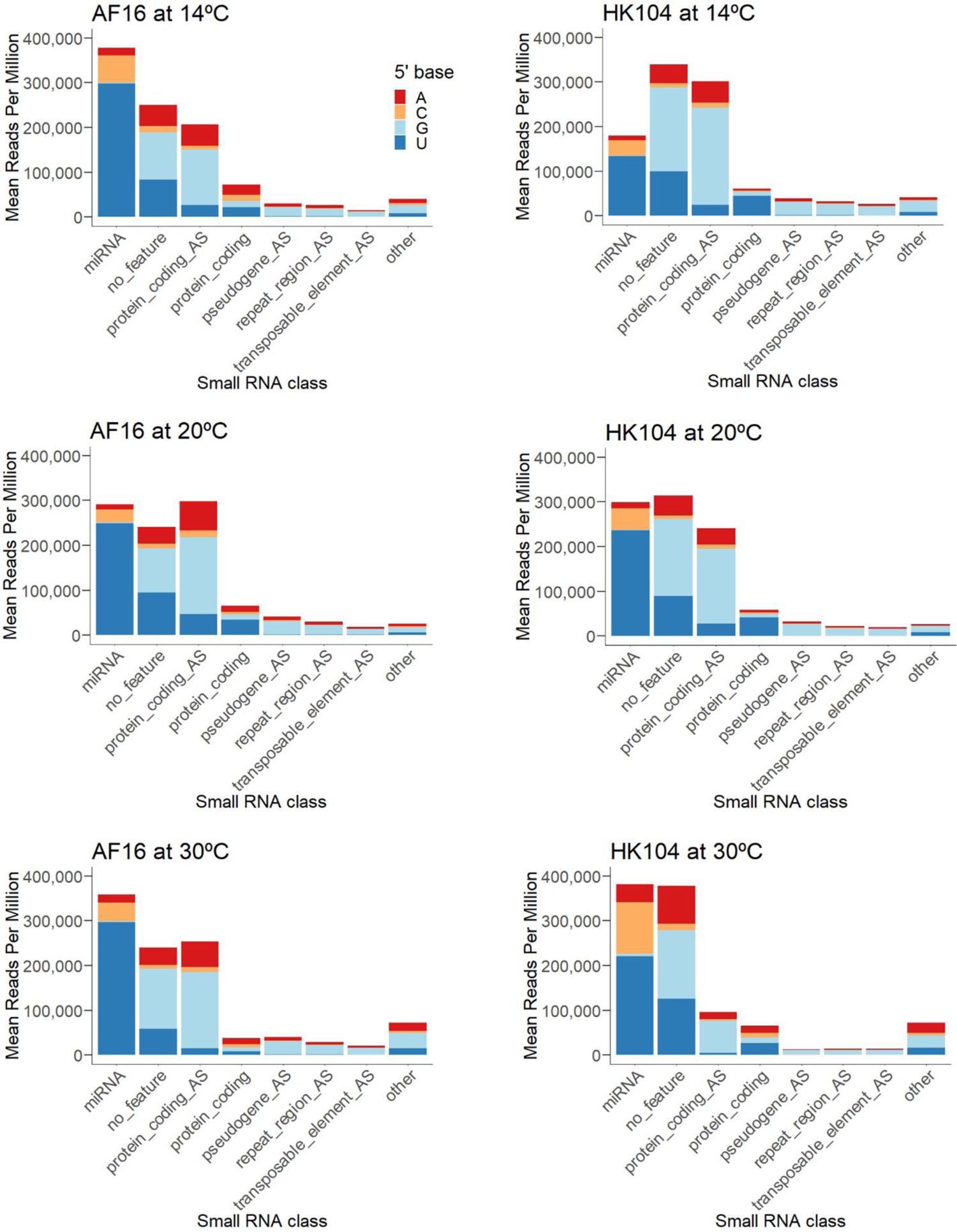
Mean expression of small RNAs mapping to different classes of genomic features in the *C. briggsae* genome, separated by the 5’ nucleotide of mapped small RNAs. “AS” indicates reads mapping antisense to a feature.

**Figure S2.**
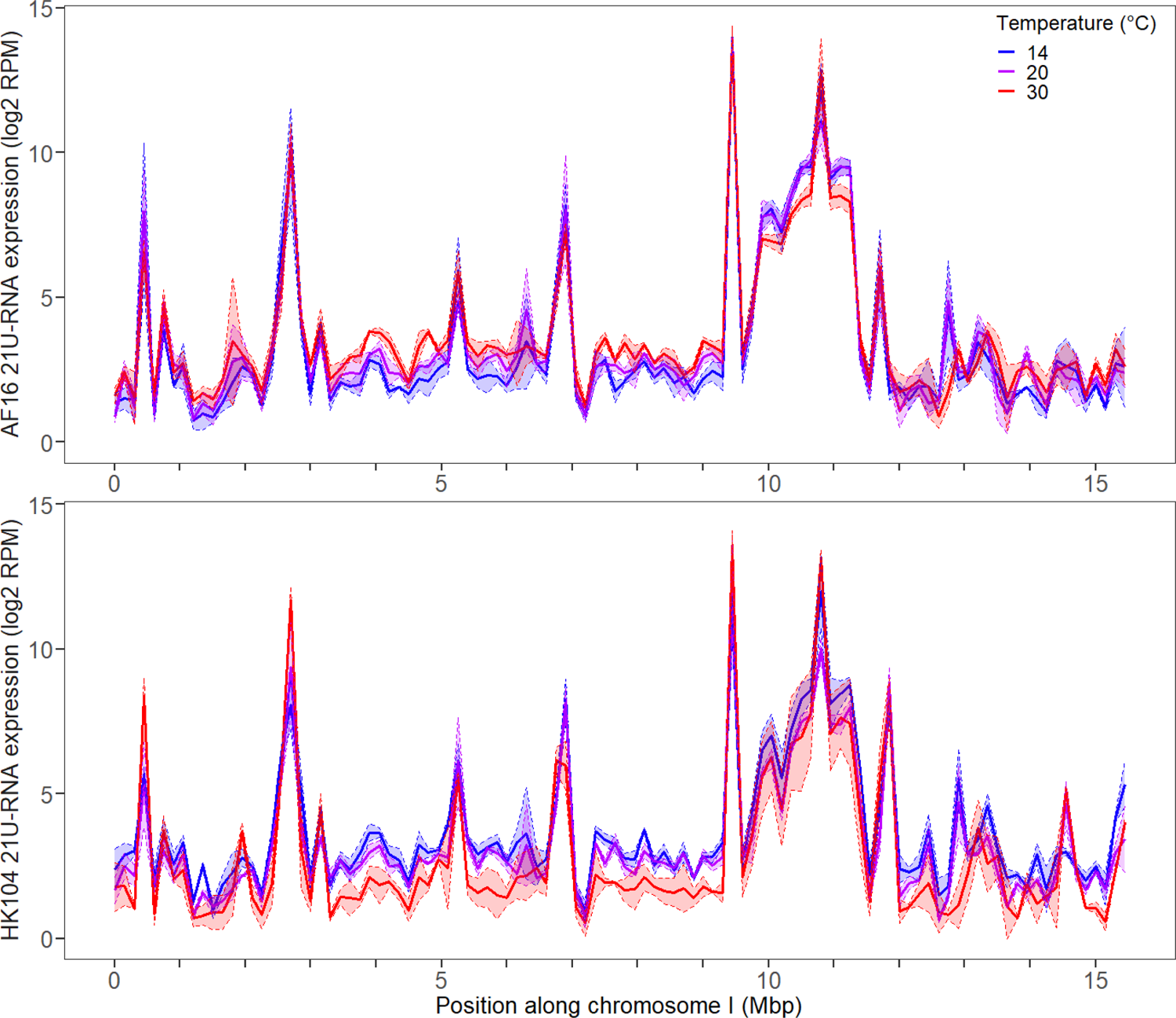
Total expression of 21U-RNAs along the length of chromosome I using genomic intervals 150kb wide, in AF16 (top) and HK104 (bottom) worms. Expression is in units of log2 Reads Per Million. Shaded regions give ±1 standard deviation around the mean for each set of replicates.

**Figure S3.**
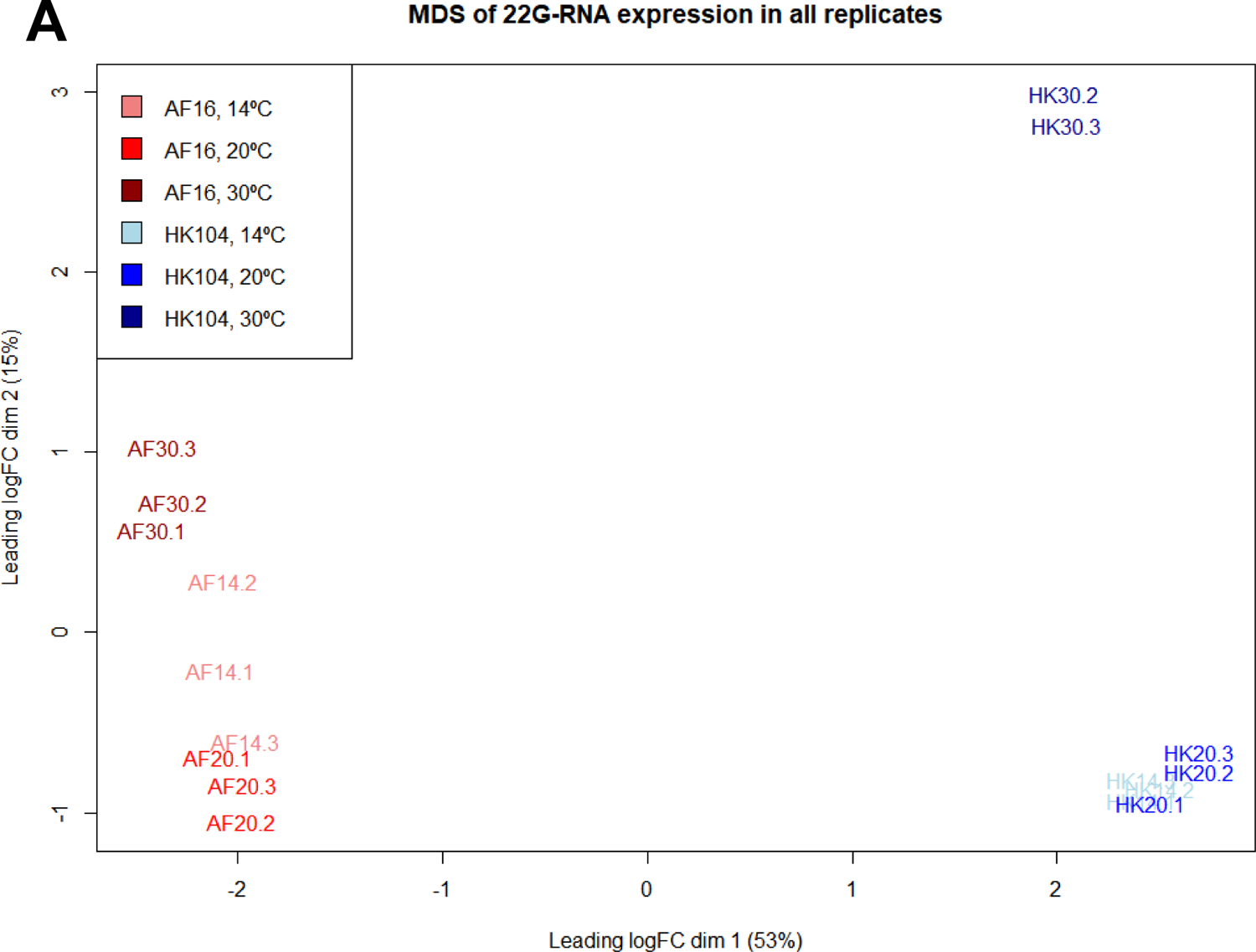
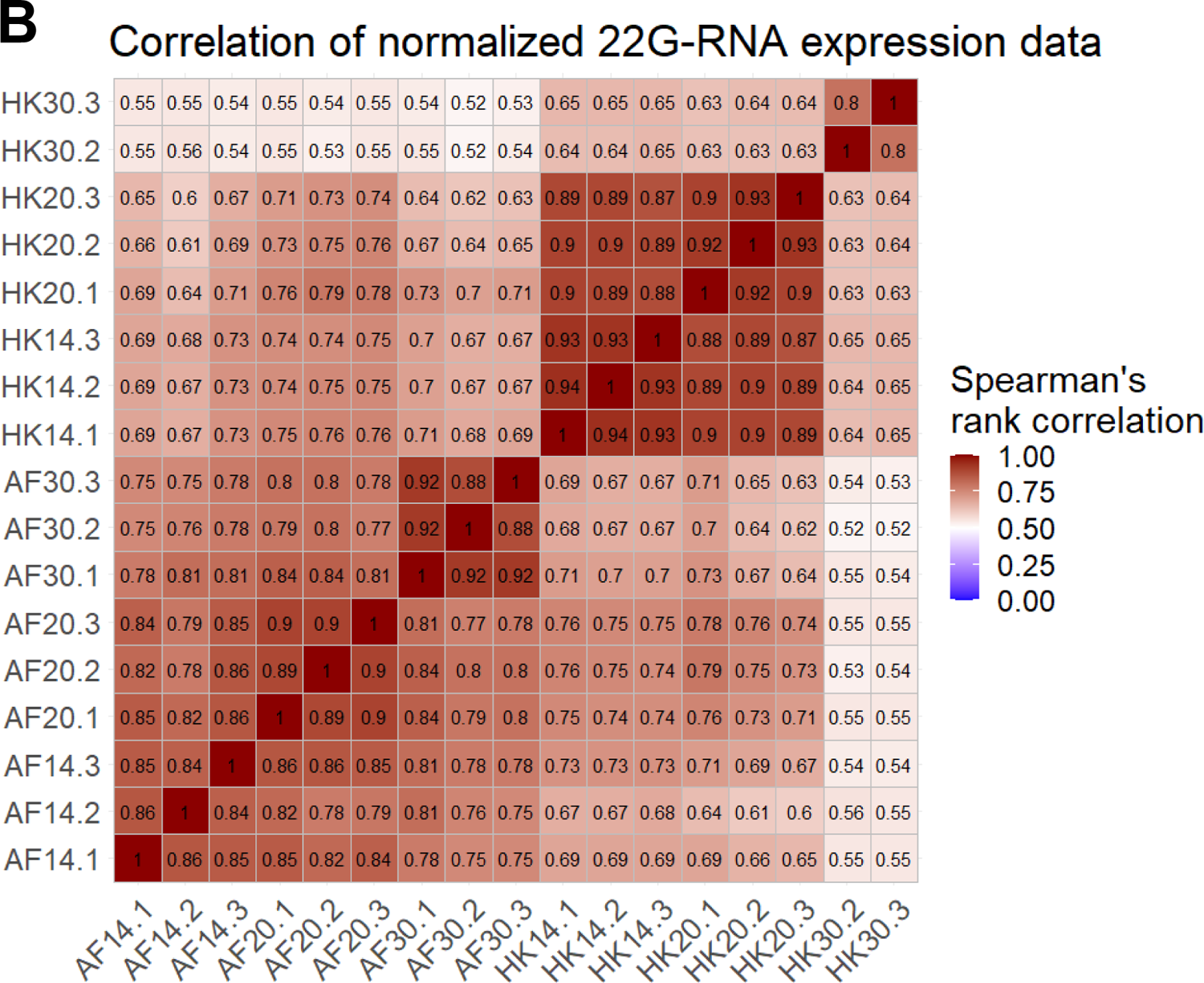
Multidimensional scaling plot (A) and correlation heatmap (B) of the 22G-RNA expression profiles of 14,283 *C. briggsae* genomic features across all 17 samples. Expression values were normalized to units of log_2_ Reads Per Million. Correlations were calculated using Spearman’s rank correlation.

**Figure S4.**
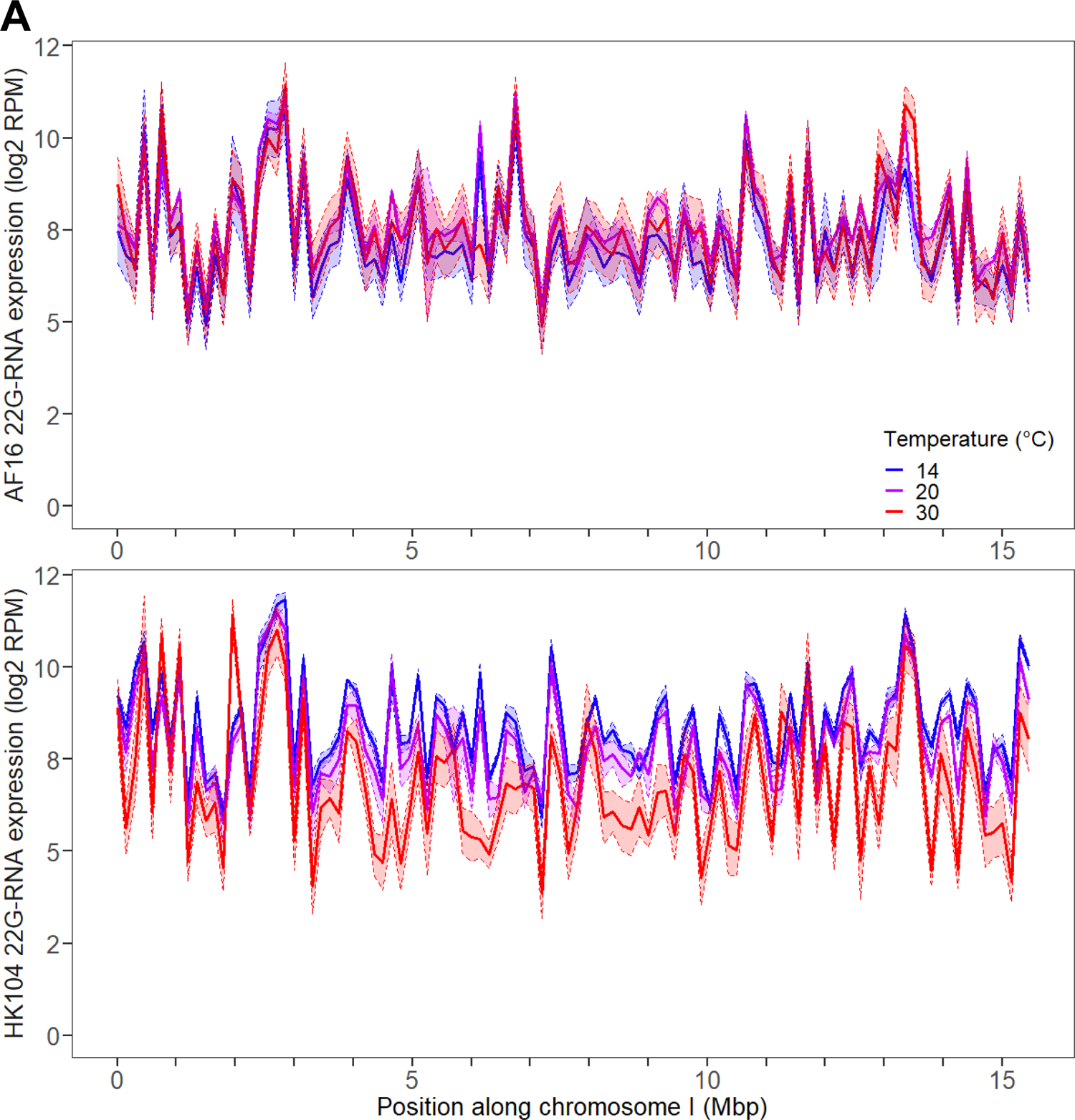
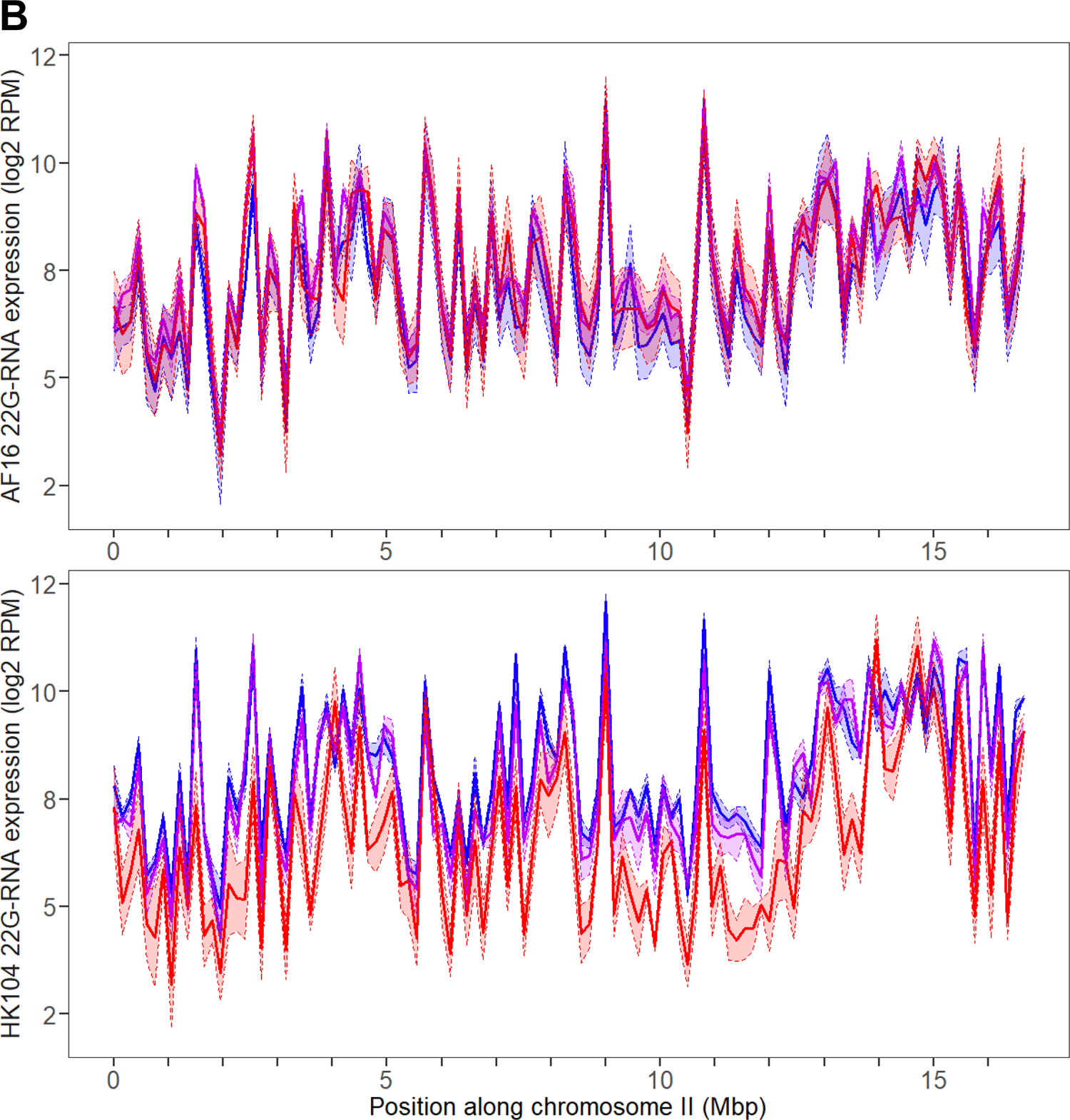
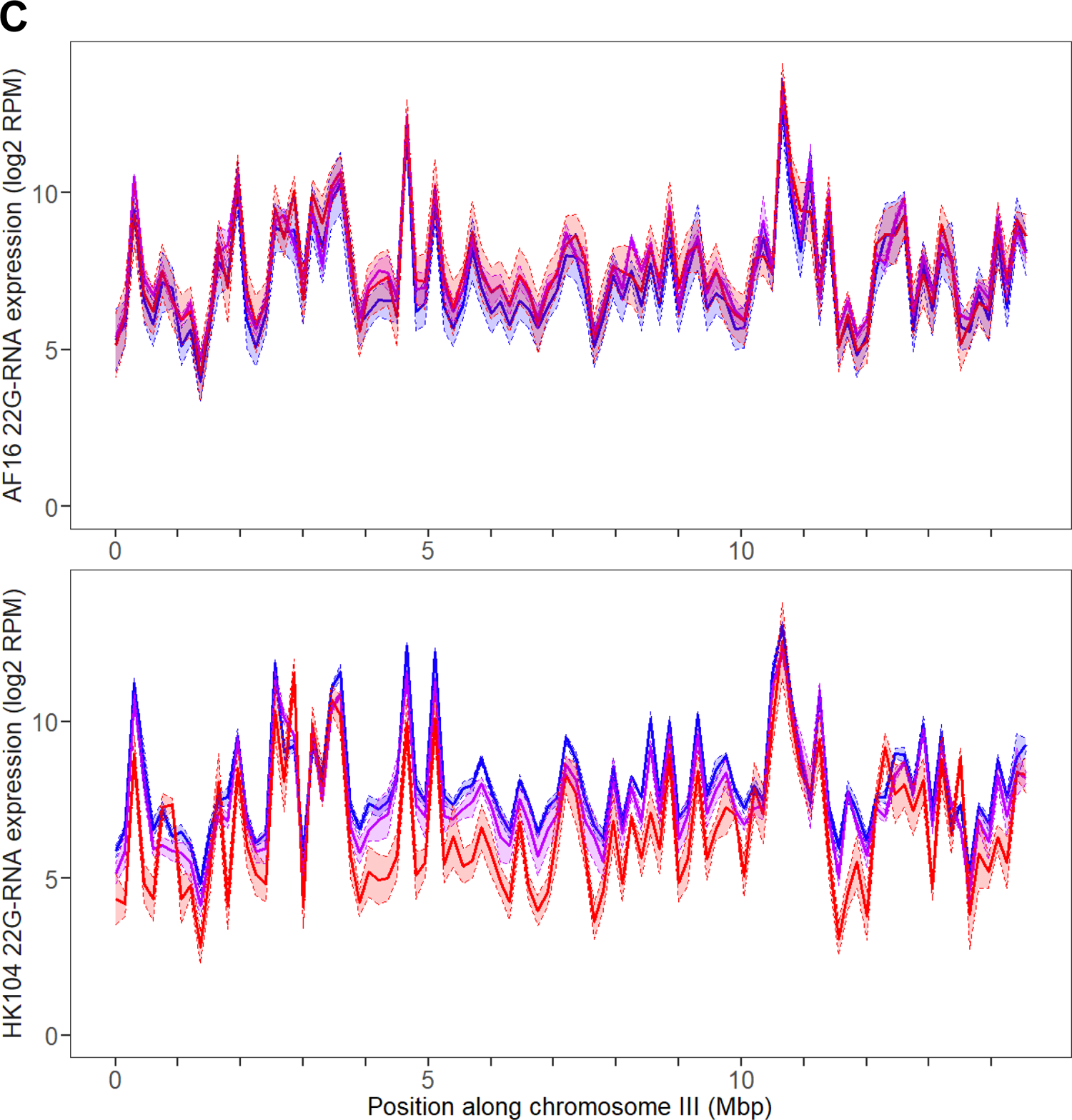
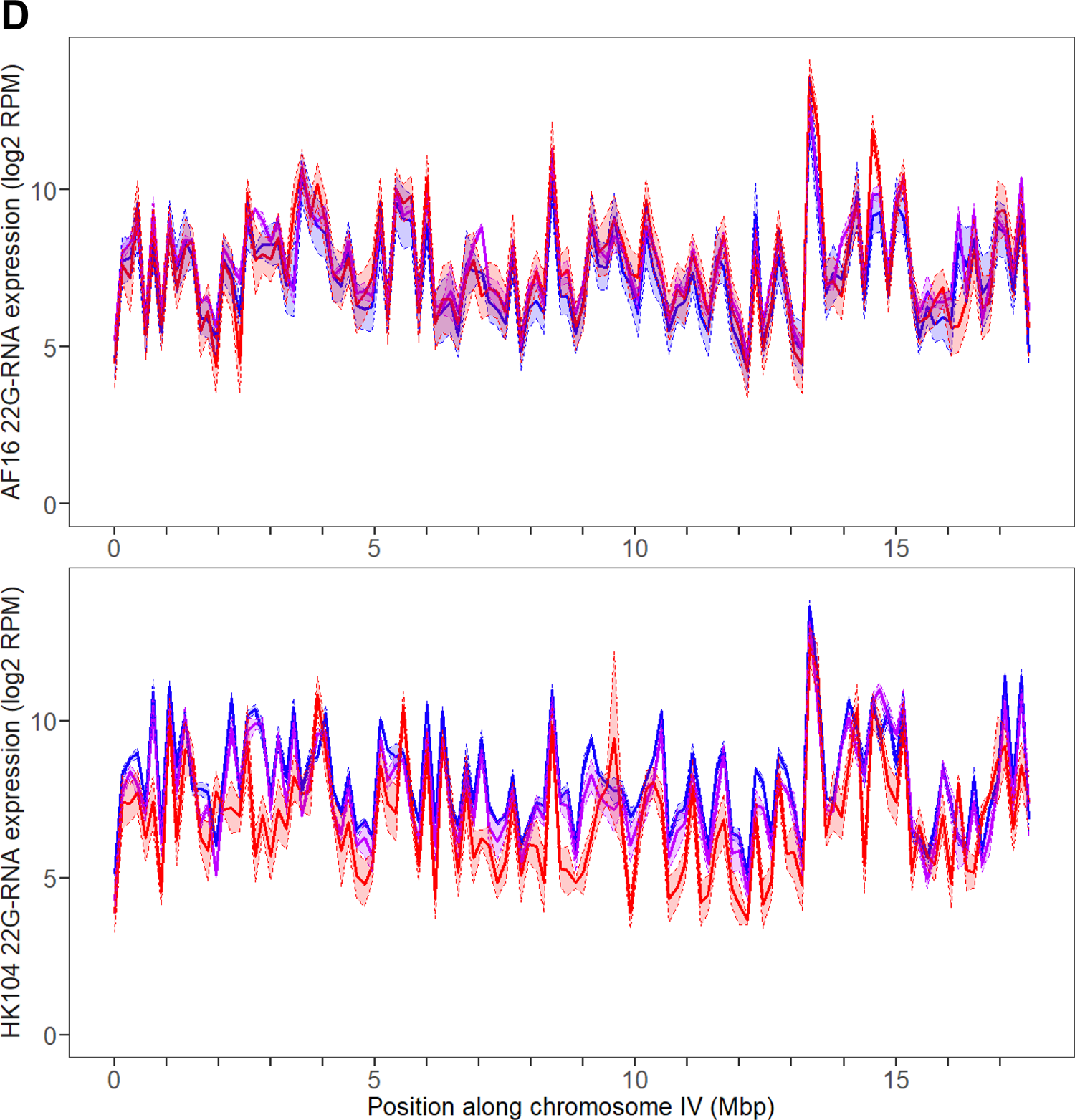
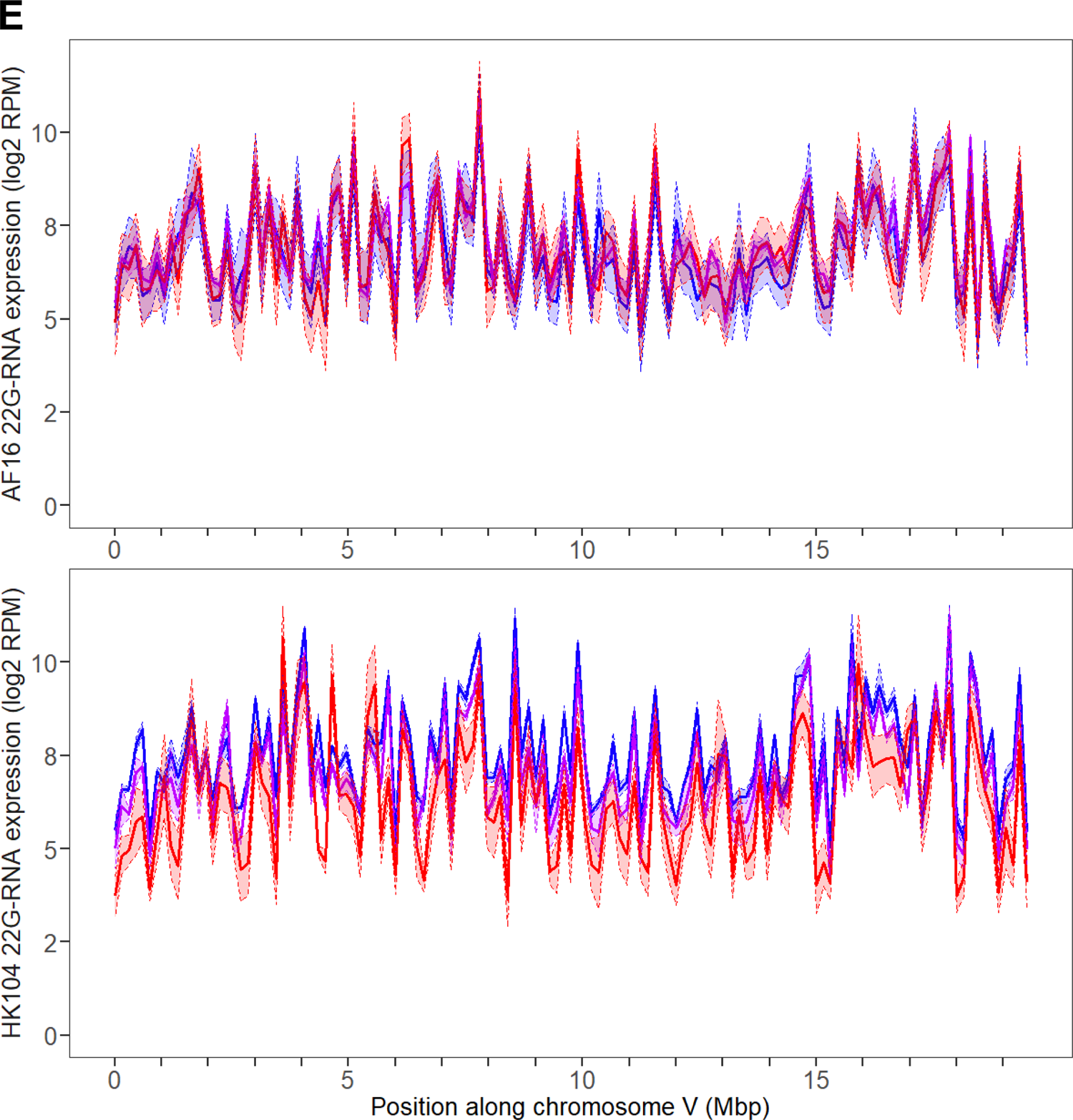
Expression of aligned 22G-RNAs across the full length of chromosomes I (A), II (B), III (C), IV (D), and V (E) in AF16 (top) and HK104 (bottom) worms, at each temperature. Expression values were averaged between replicates, using genomic intervals 150 kilobases wide. Expression is in units of log2 Reads Per Million (RPM). Shaded areas extend ±1 standard deviation around the mean expression of each interval.

**Figure S5.**
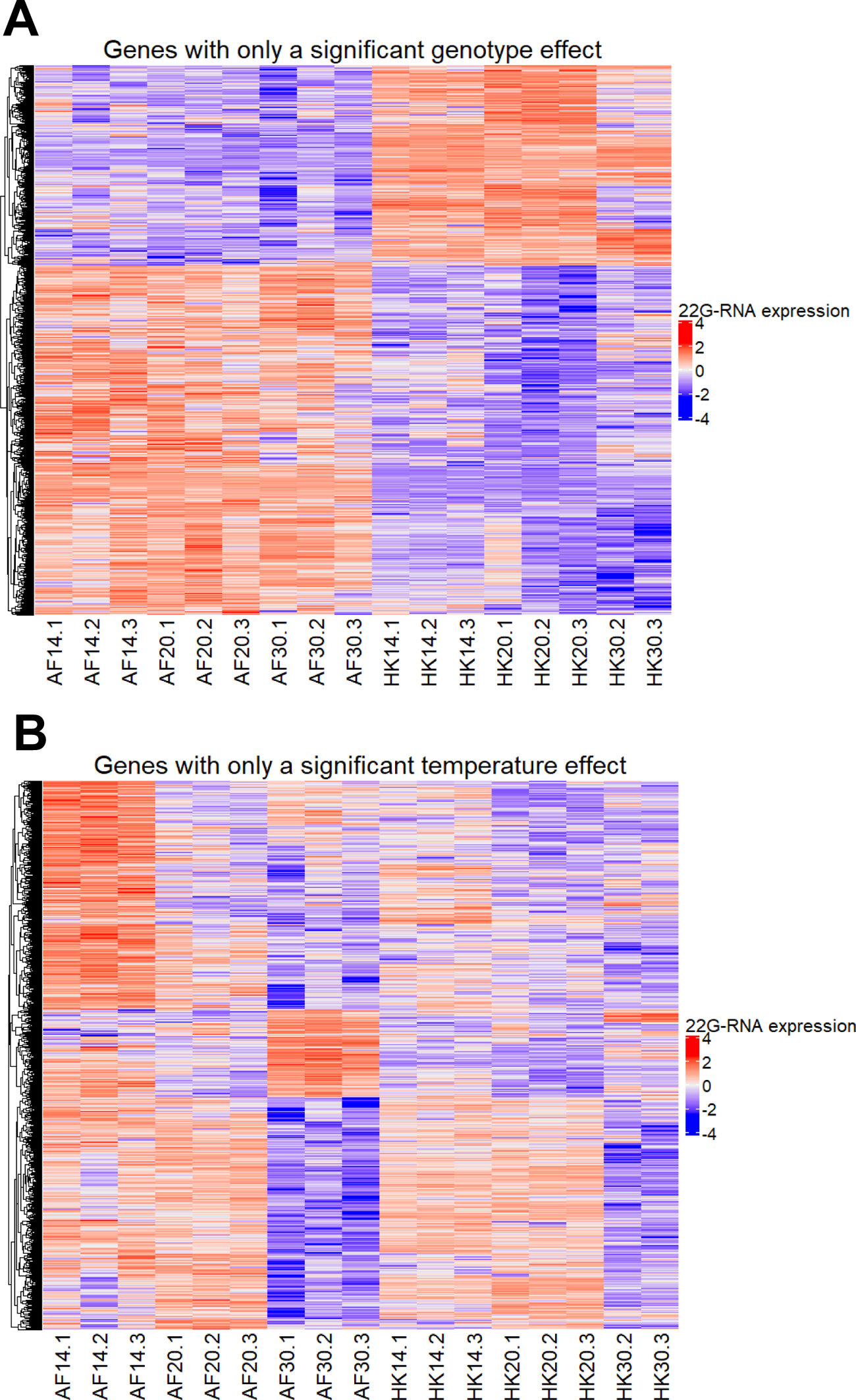
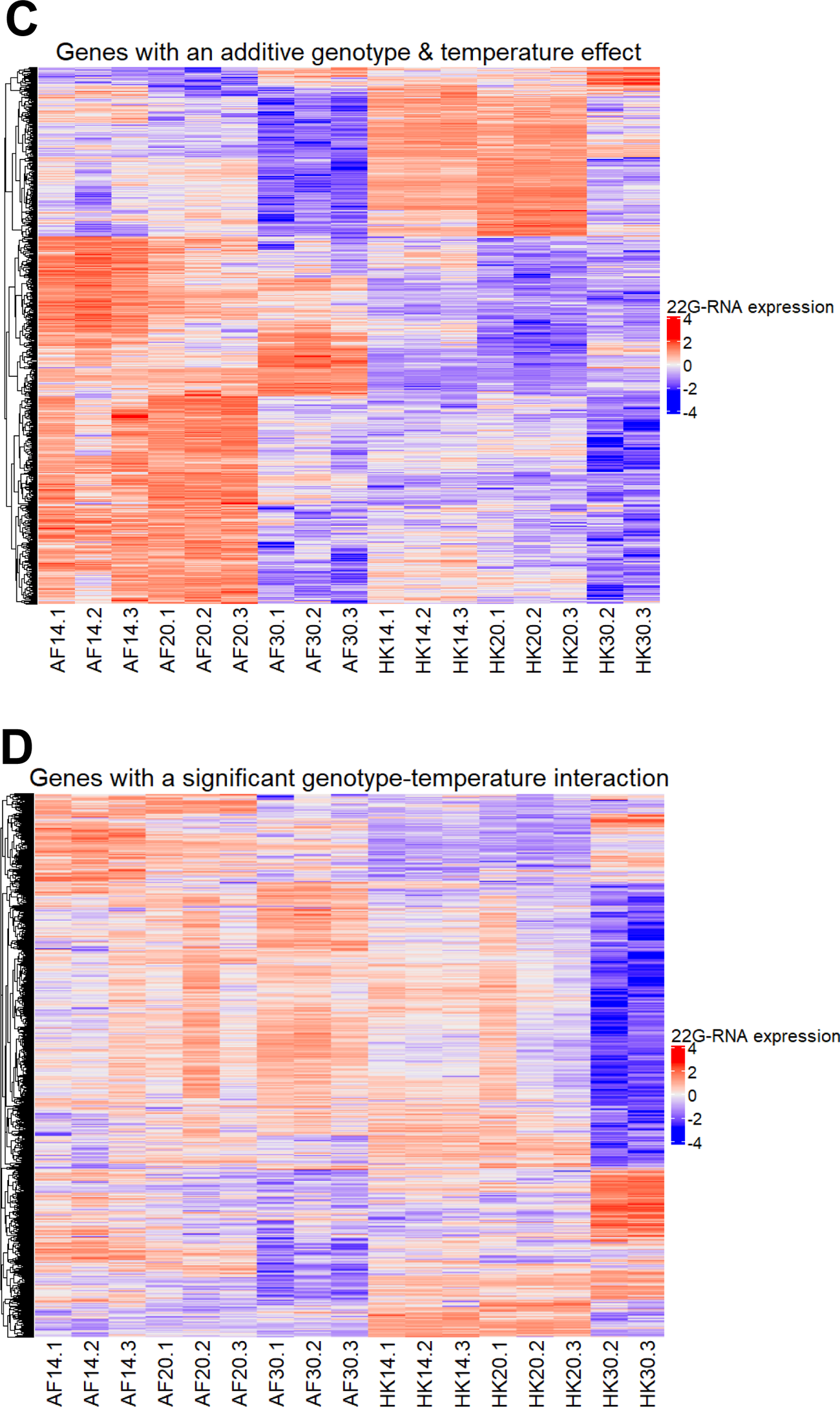
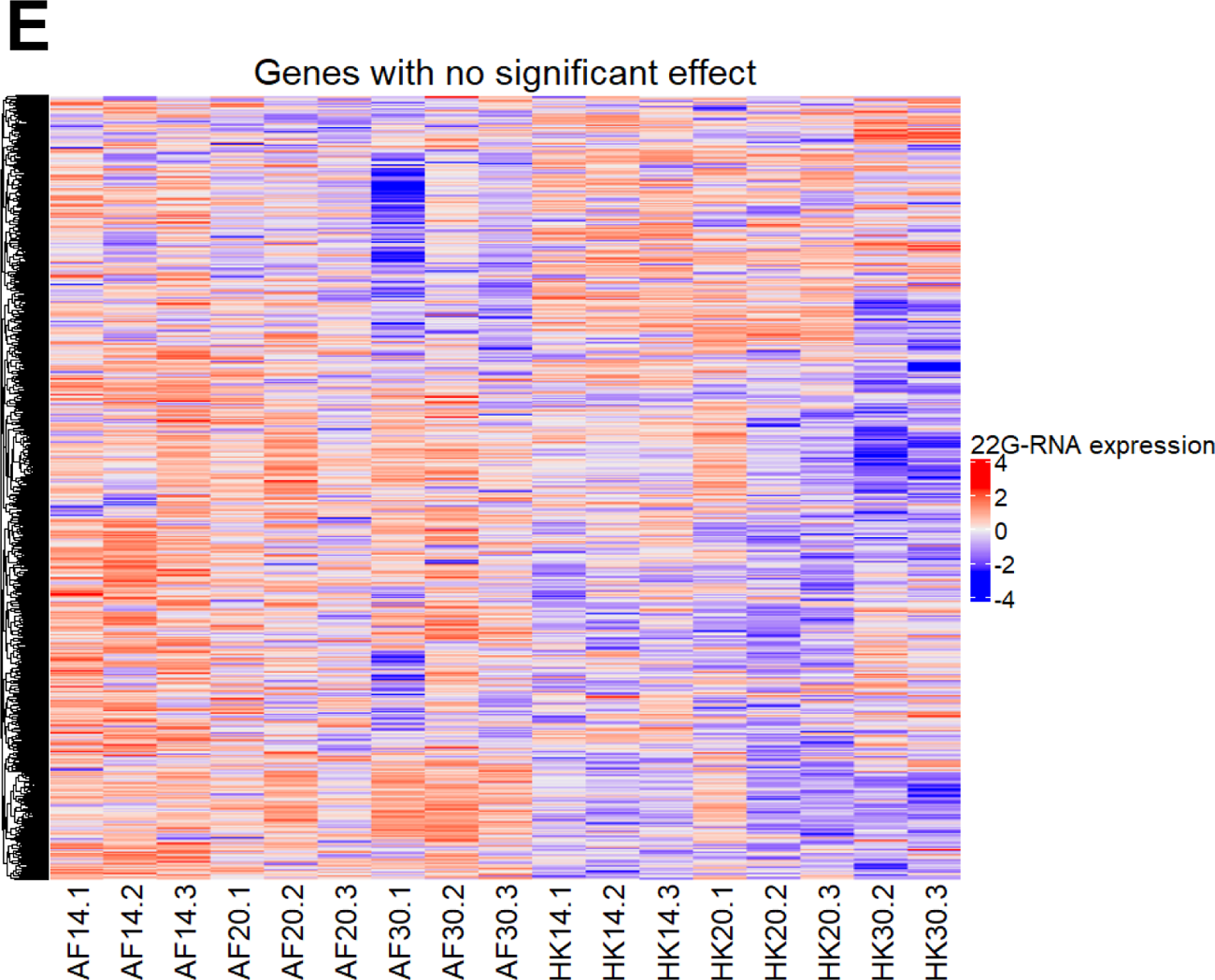
22G-RNA expression profiles across all 17 samples, for each of the 5 categories of genomic features defined based on 22G-RNA expression effects (A – E). Features include protein-coding genes, pseudogenes, repeats, and transposons. Expression values for each feature are scaled using row normalization. Rows in each heatmap are ordered based on hierarchical clustering.

**Figure S6.**
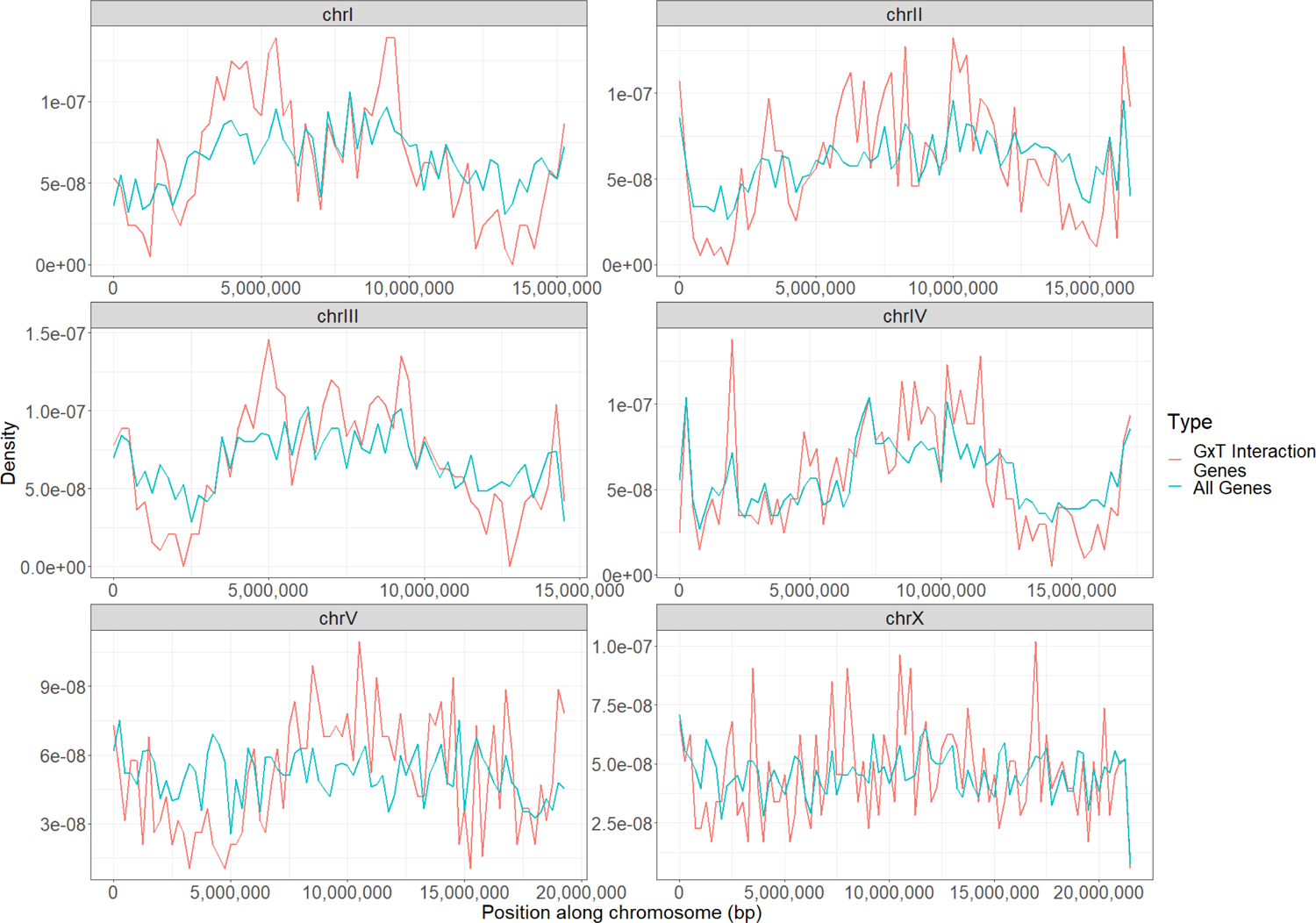
Distribution of protein-coding genes showing a genotype-temperature interaction on 22G-RNA expression across the length of each *C. briggsae* chromosome, compared to the genomic distribution of all *C. briggsae* protein-coding genes.

**Figure S7.**
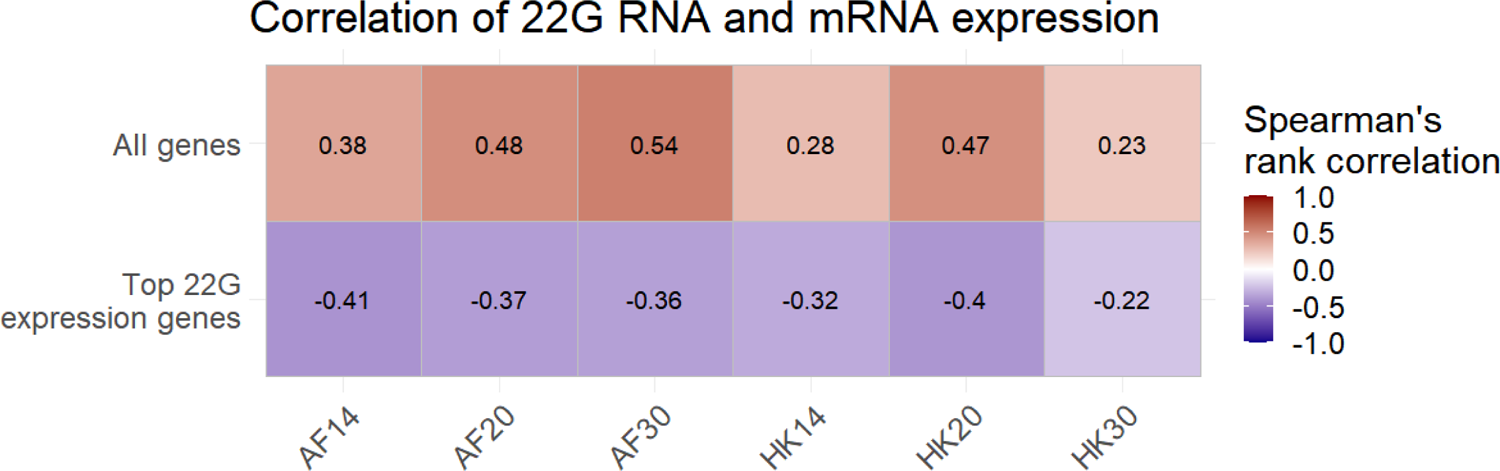
Correlation between 22G-RNA and mRNA expression for all 6 genotype-temperature combinations. Correlations were calculated using the set of all 12,623 genes (top row), as well as using only the genes in each genotype-temperature combination that were in the top 10% of 22G-RNA expression (bottom row). Expression values were normalized to units of log_2_ Reads Per Million. Correlations were calculated using Spearman’s rank correlation.

**Figure S8.**
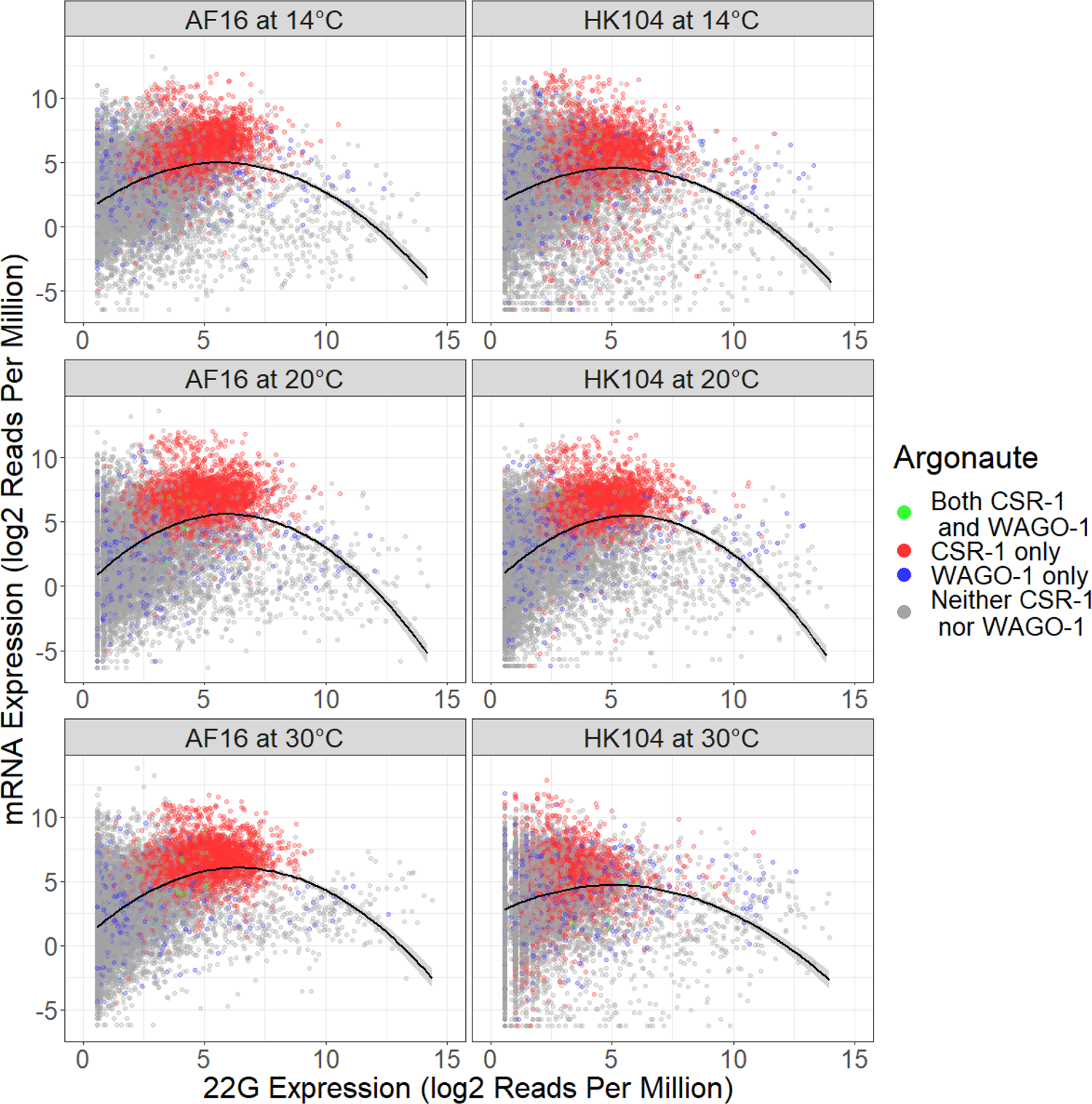
22G-RNA and mRNA expression for each genotype-temperature combination, for 12,623 genes with 22G-RNA and mRNA expression data available. A quadratic curve of best fit is also shown, with the shaded area around the curve representing the standard error. Colors indicate whether each gene is a target of CSR-1, WAGO-1, both, or neither.

**Figure S9.**
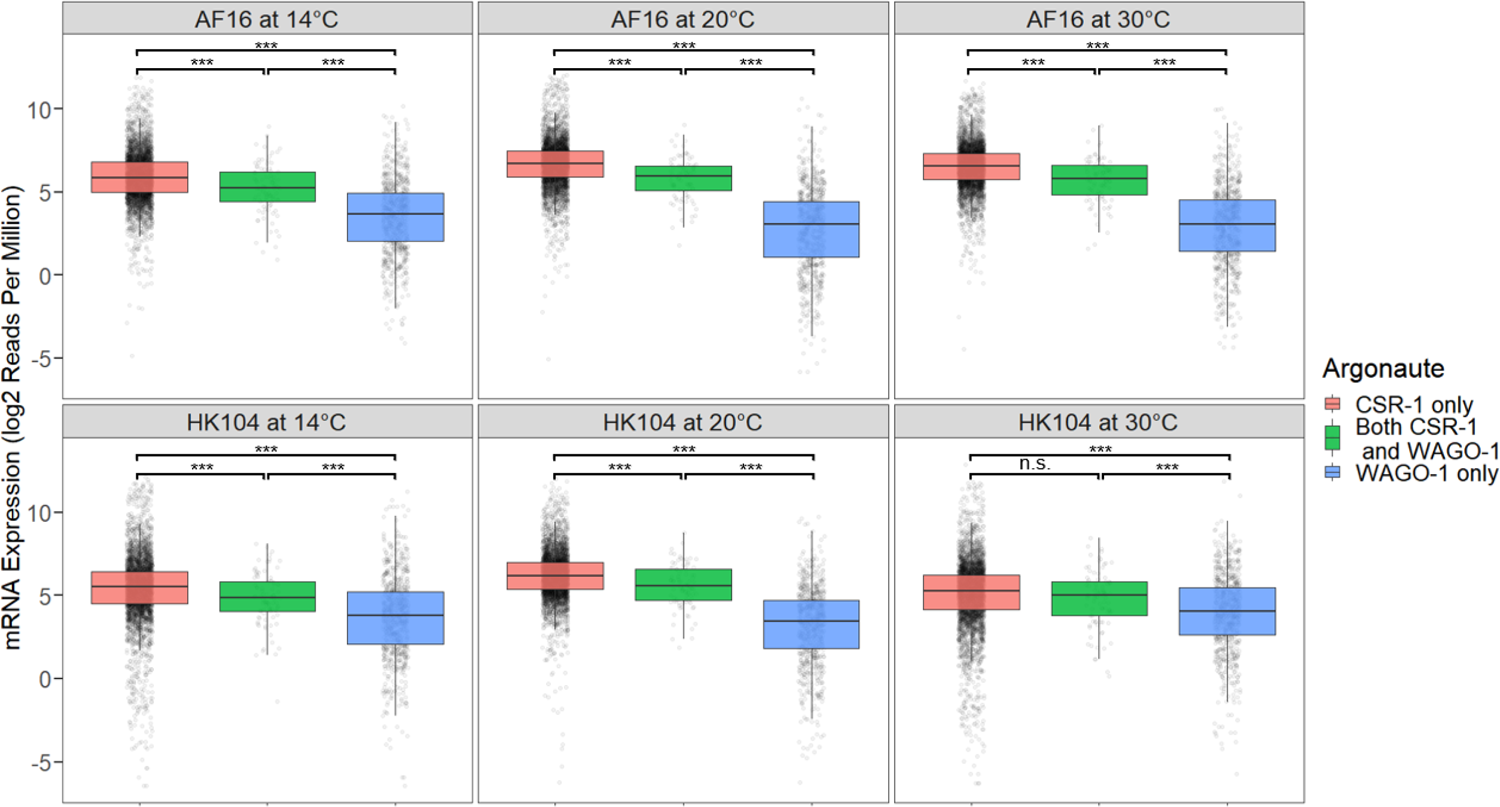
Distribution of mRNA expression values in each genotype-temperature combination, for genes that are targeted by CSR-1 but not WAGO-1 (3804 genes), both CSR-1 and WAGO-1 simultaneously (87 genes), or WAGO-1 but not CSR-1 (749 genes). Significant differences between sets of genes were determined using a Wilcoxon rank sum test, with *** indicating *p* < 0.001 and n.s. indicating non-significance.

**Figure S10.**
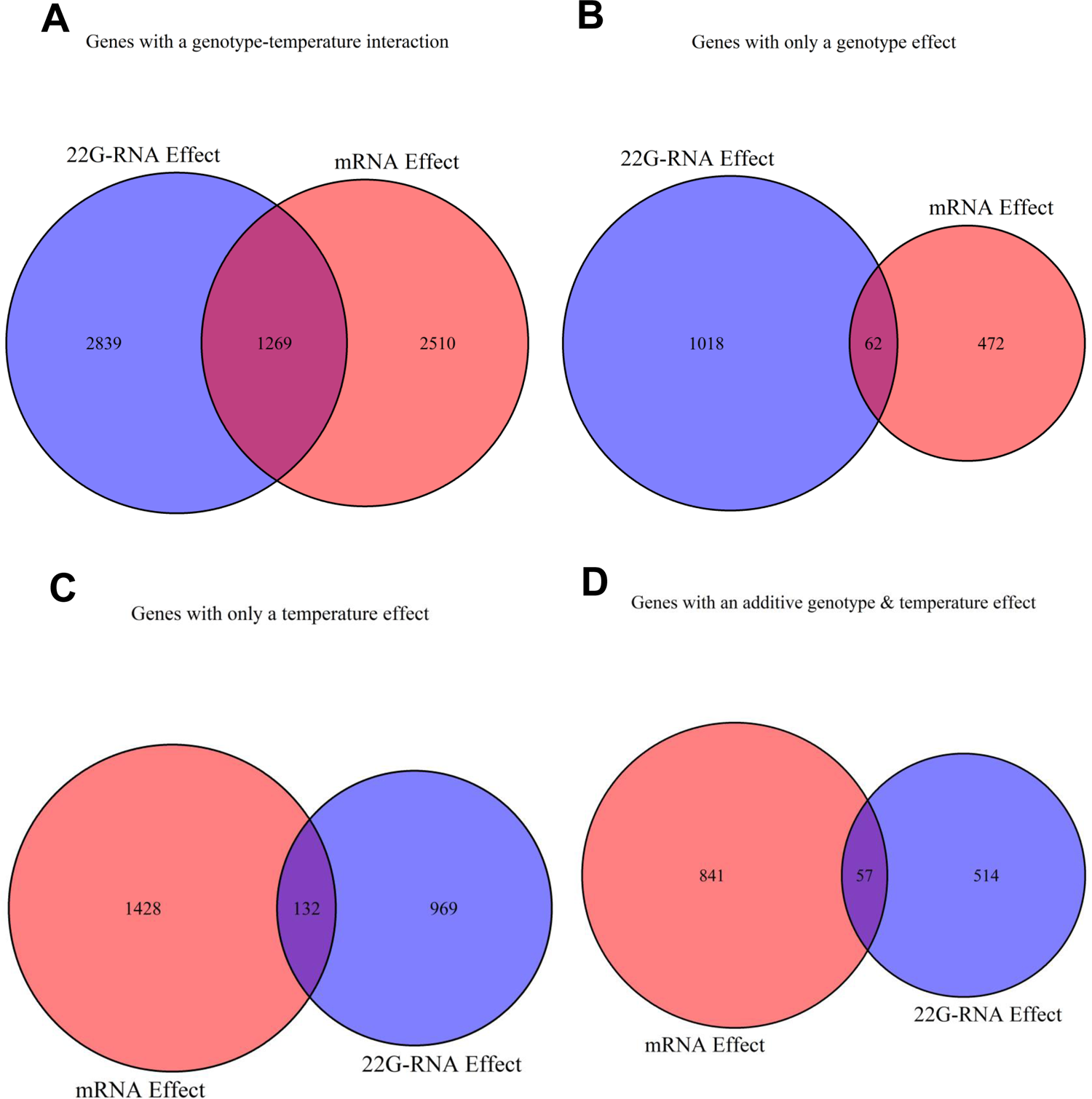
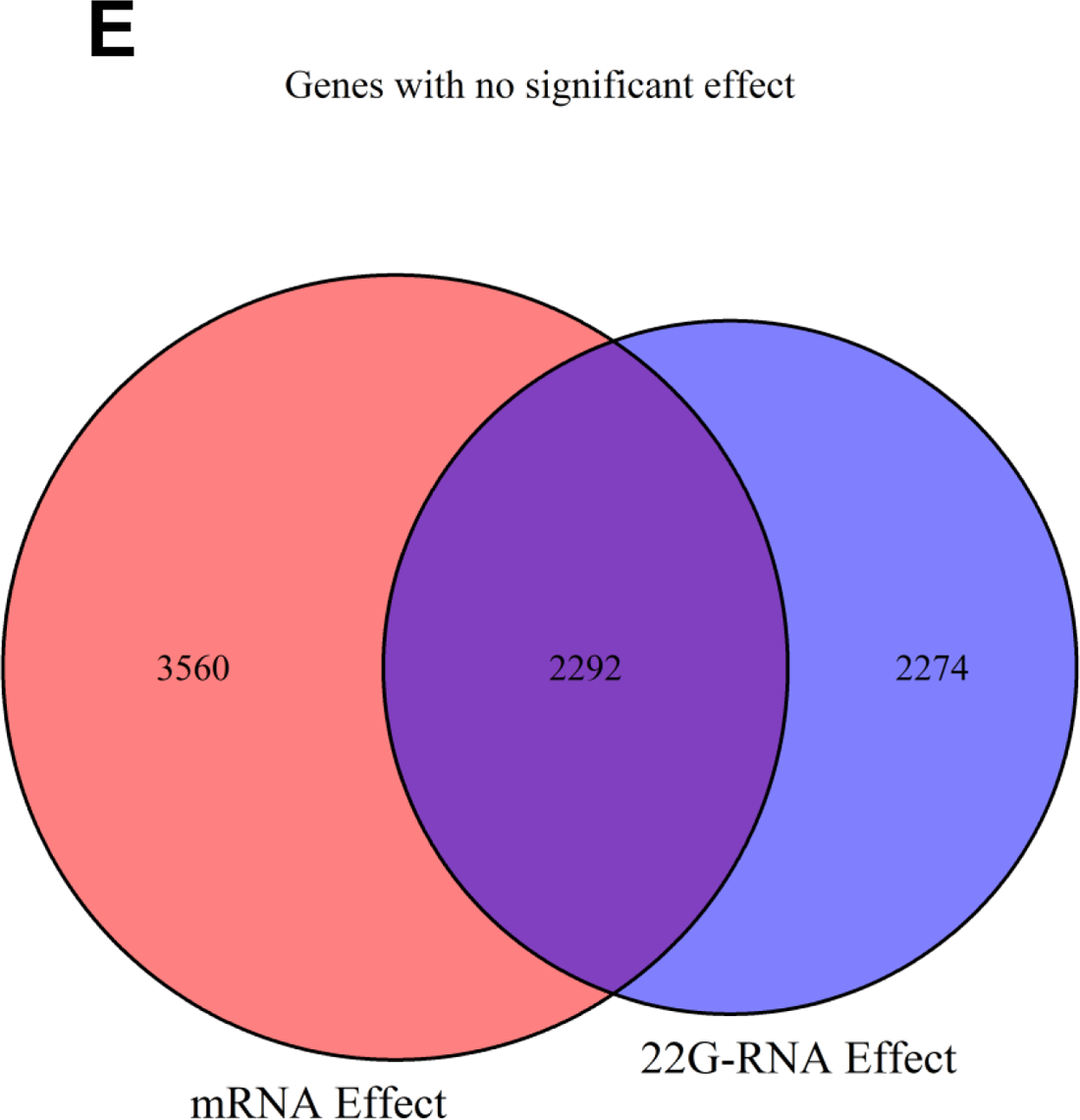
Overlap between genes showing a given effect on 22G-RNA expression, and genes showing the same effect on mRNA expression, for different categories of expression effects (A – E).

**Figure S11.**
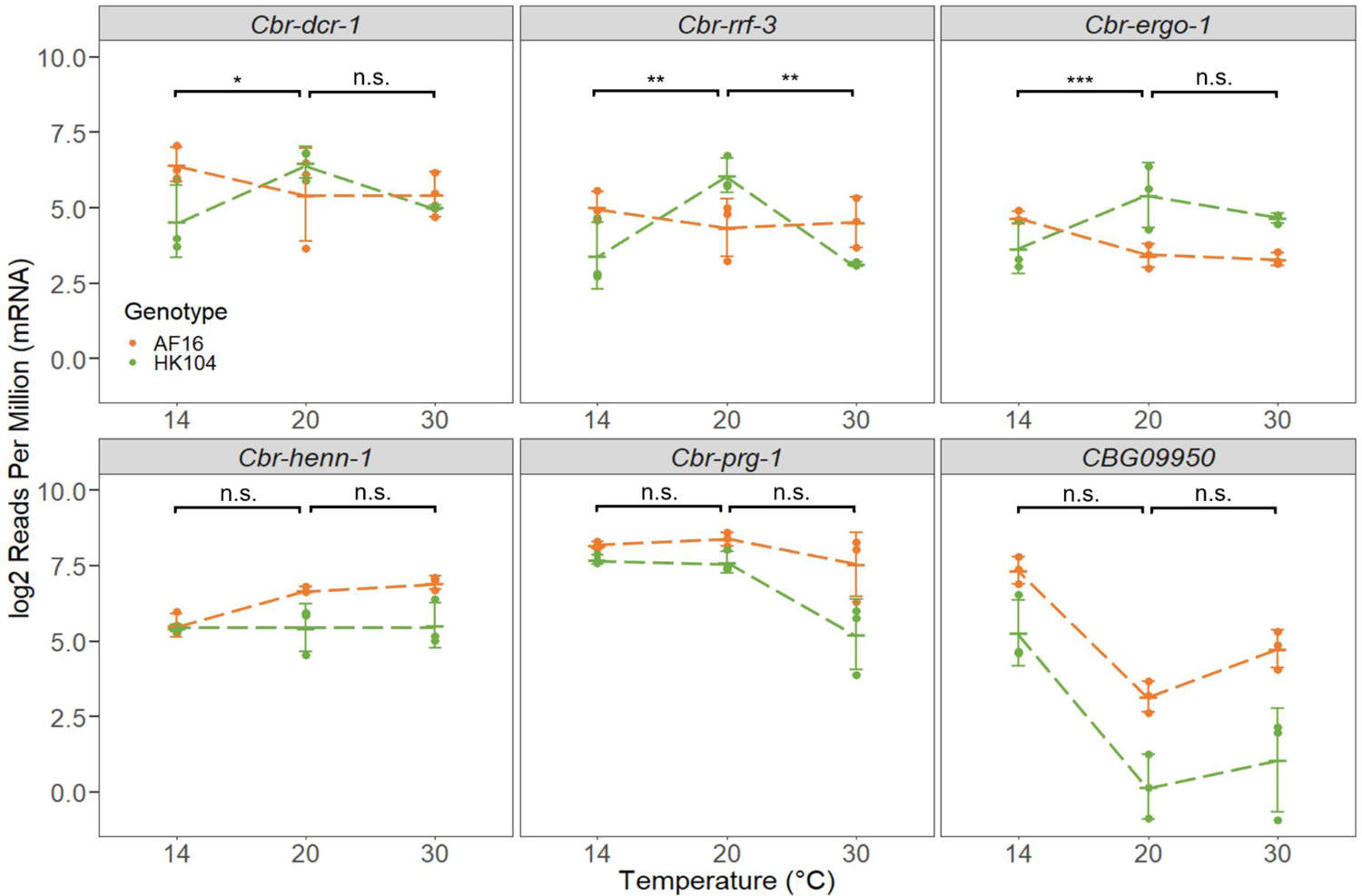
mRNA expression of different genes involved in the 26G-RNA and piRNA pathways, across genotypes and temperatures. *CBG09950* is the *C. briggsae* ortholog of both *alg-3* and *alg-4*. Error bars give ±1 standard deviation around the mean for each set of replicates. Dashed lines connect mean expression values for each genotype across temperatures. Linear modeling was used to determine the significance of genotype-temperature interactions between 14°C and 20°C, and between 20°C and 30°C, with * indicating *p* < 0.05, ** indicating *p* < 0.01, *** indicating *p* < 0.001, and n.s. indicating non-significance.

**Figure S12.**
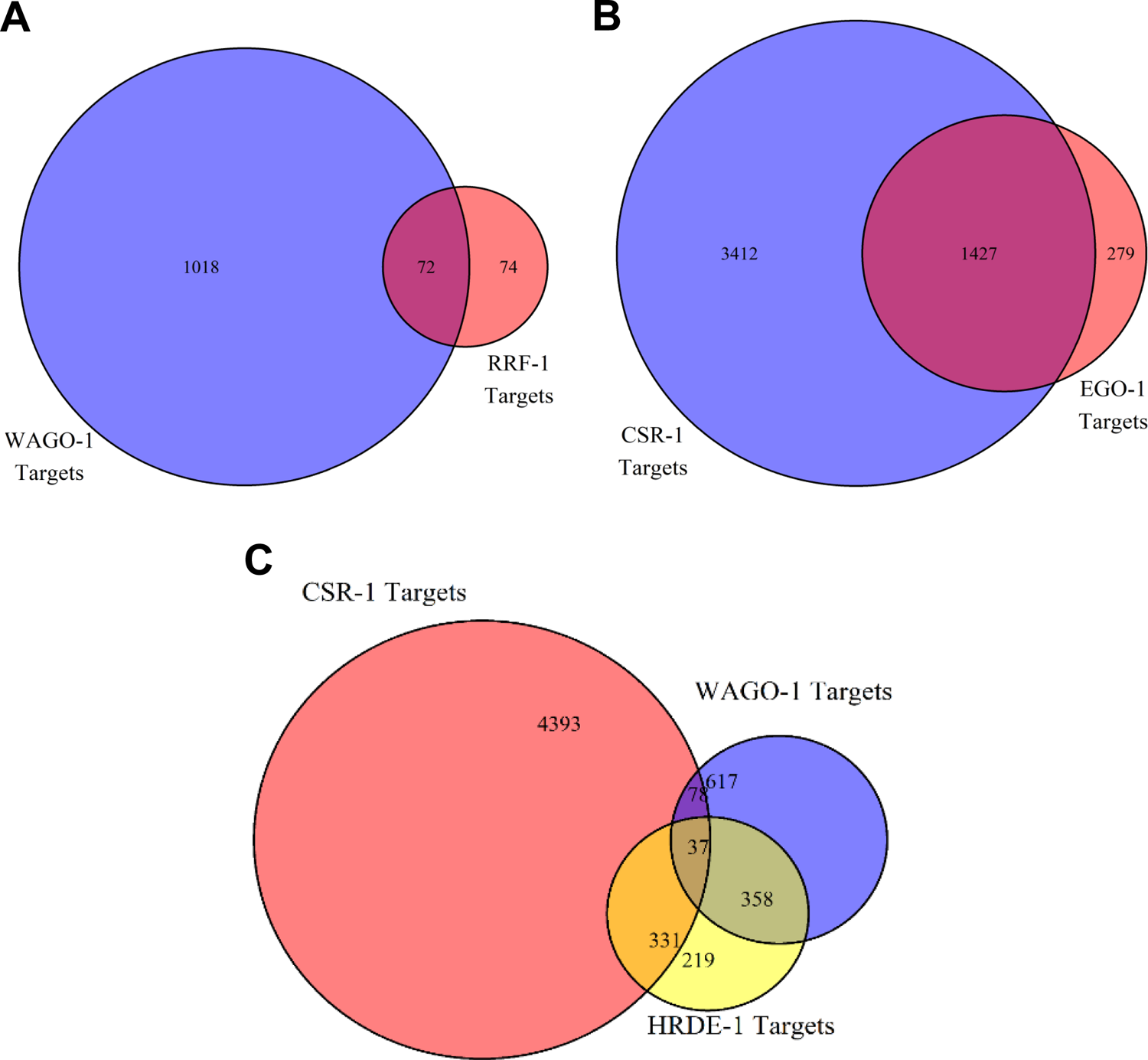
Overlap between the genes targeted by WAGO-1 or RRF-1 (A), between the genes targeted by CSR-1 or EGO-1 (B), and between the genes targeted by CSR-1, WAGO-1, or HRDE-1 (C).

**Figure S13.**
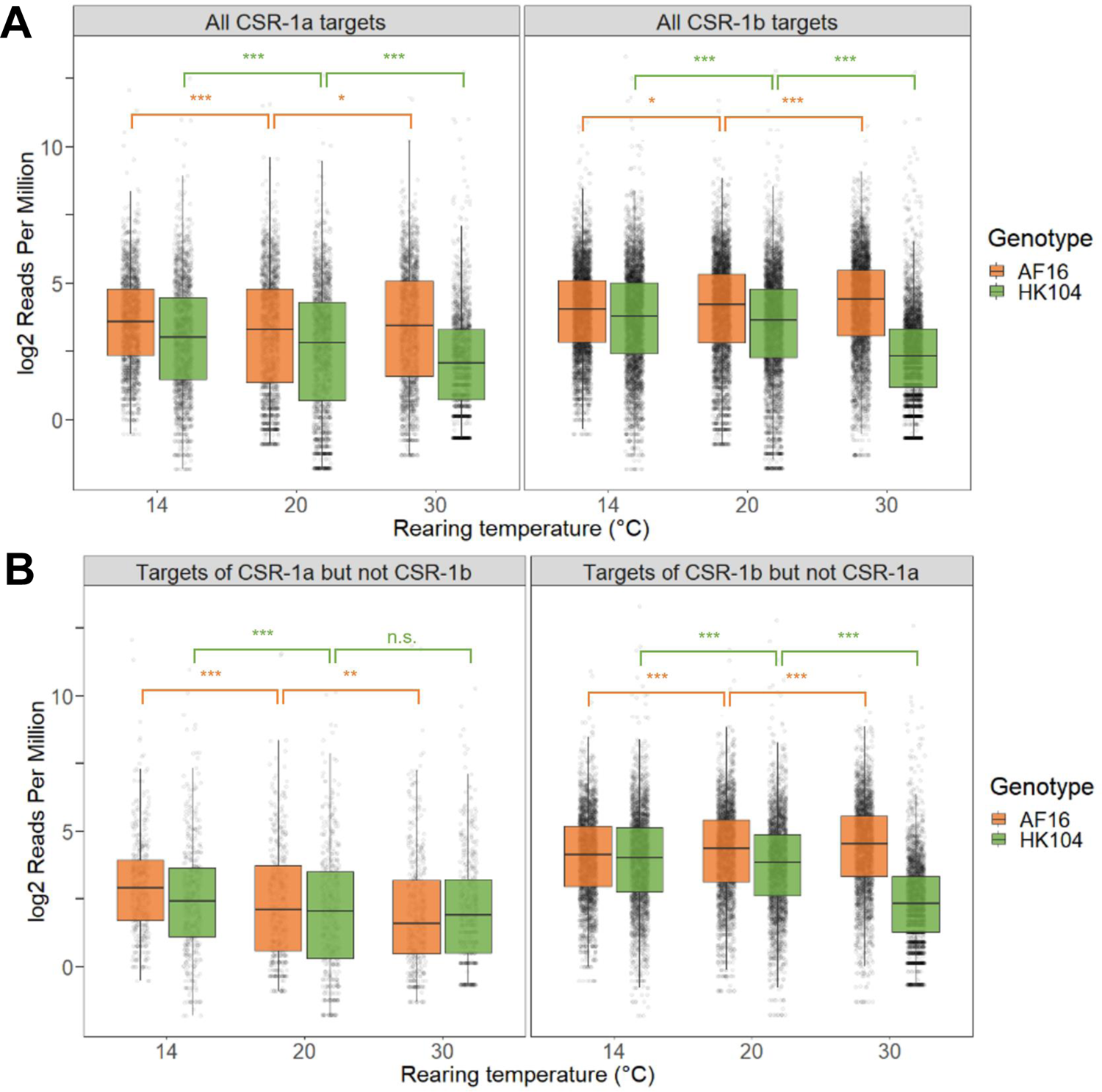
22G-RNA expression of all genes targeted by CSR-1a or CSR-1b (A) as well as genes exclusively targeted by only one CSR-1 isoform and not the other (B), for each genotype-temperature combination. Significant differences between temperature treatments of the same genotype were determined using a Wilcoxon rank sum test, with * indicating *p* < 0.05, ** indicating *p* < 0.01, *** indicating *p* < 0.001, and n.s. indicating non-significance.

**Figure S14.**
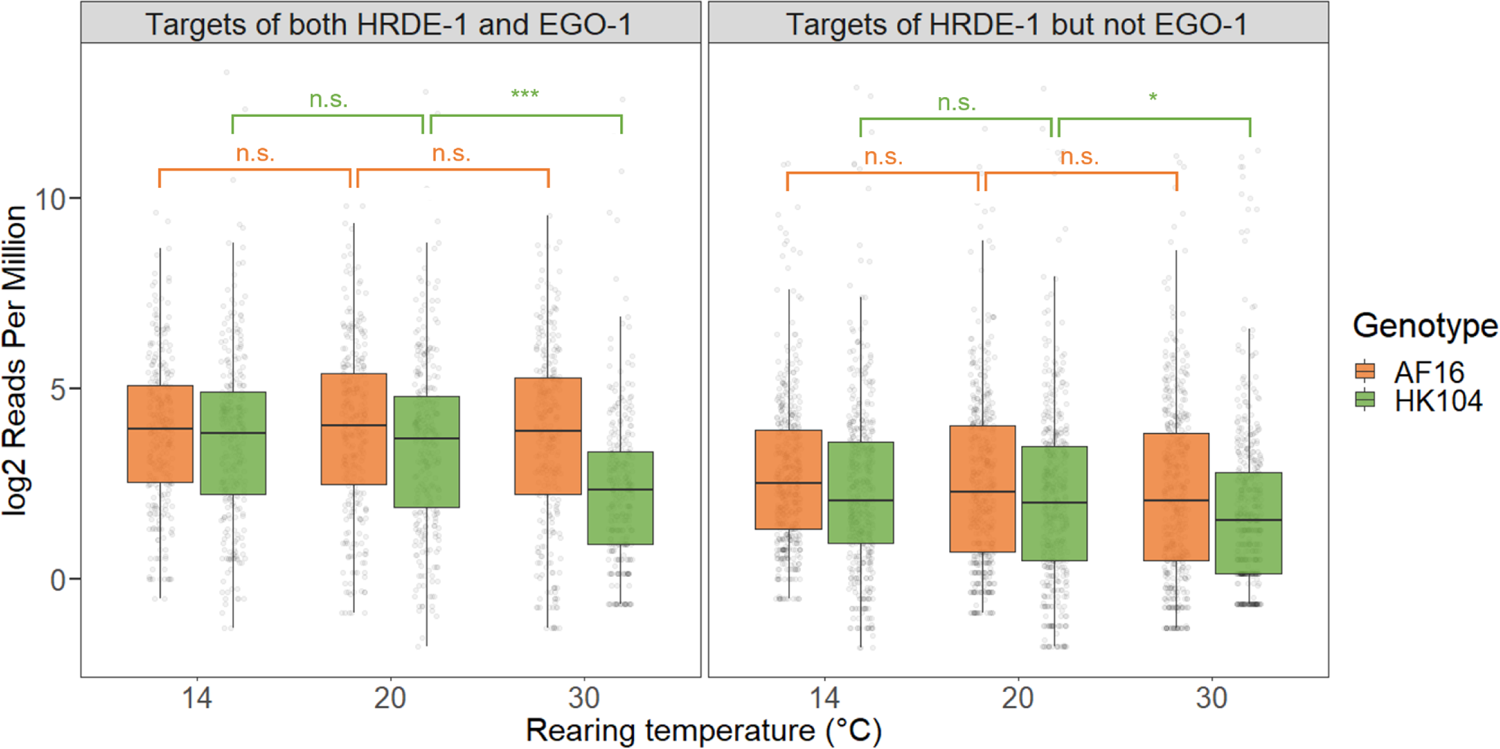
22G-RNA expression of genes targeted by HRDE-1 for each genotype-temperature combination, separated based on whether or not the gene is also targeted by EGO-1. Significant differences between temperature treatments of the same genotype were determined using a Wilcoxon rank sum test, with * indicating *p* < 0.05, *** indicating *p* < 0.001, and n.s. indicating non-significance.

**Figure S15.**
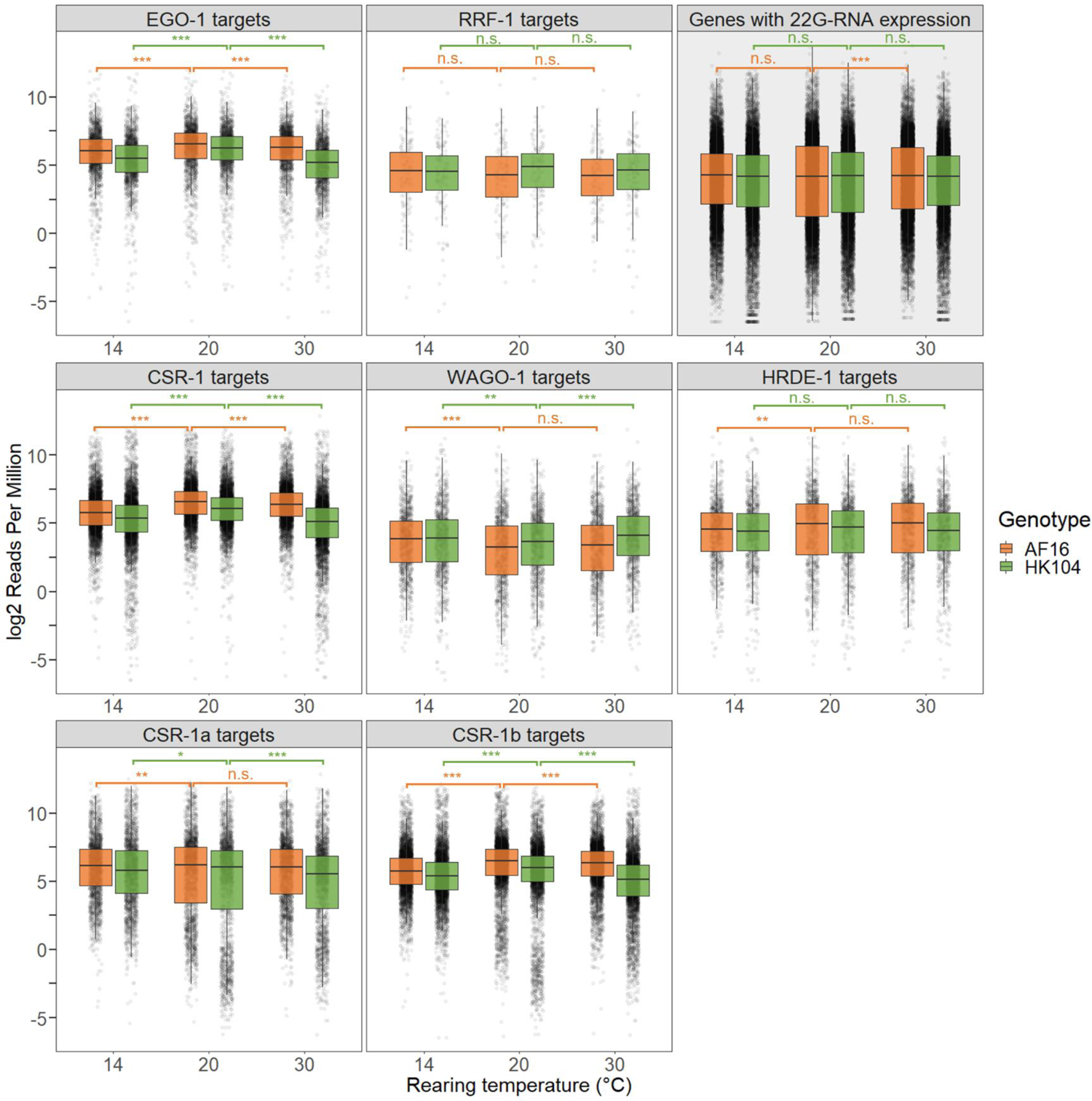
mRNA expression of genes targeted by EGO-1, RRF-1, CSR-1, WAGO-1, HRDE-1, CSR-1a, or CSR-1b, as well as all genes showing 22G-RNA expression, for each genotype-temperature combination. Significant differences between temperature treatments of the same genotype were determined using a Wilcoxon rank sum test, with * indicating *p* < 0.05, ** indicating *p* < 0.01, *** indicating *p* < 0.001, and n.s. indicating non-significance.

**Figure S16.**
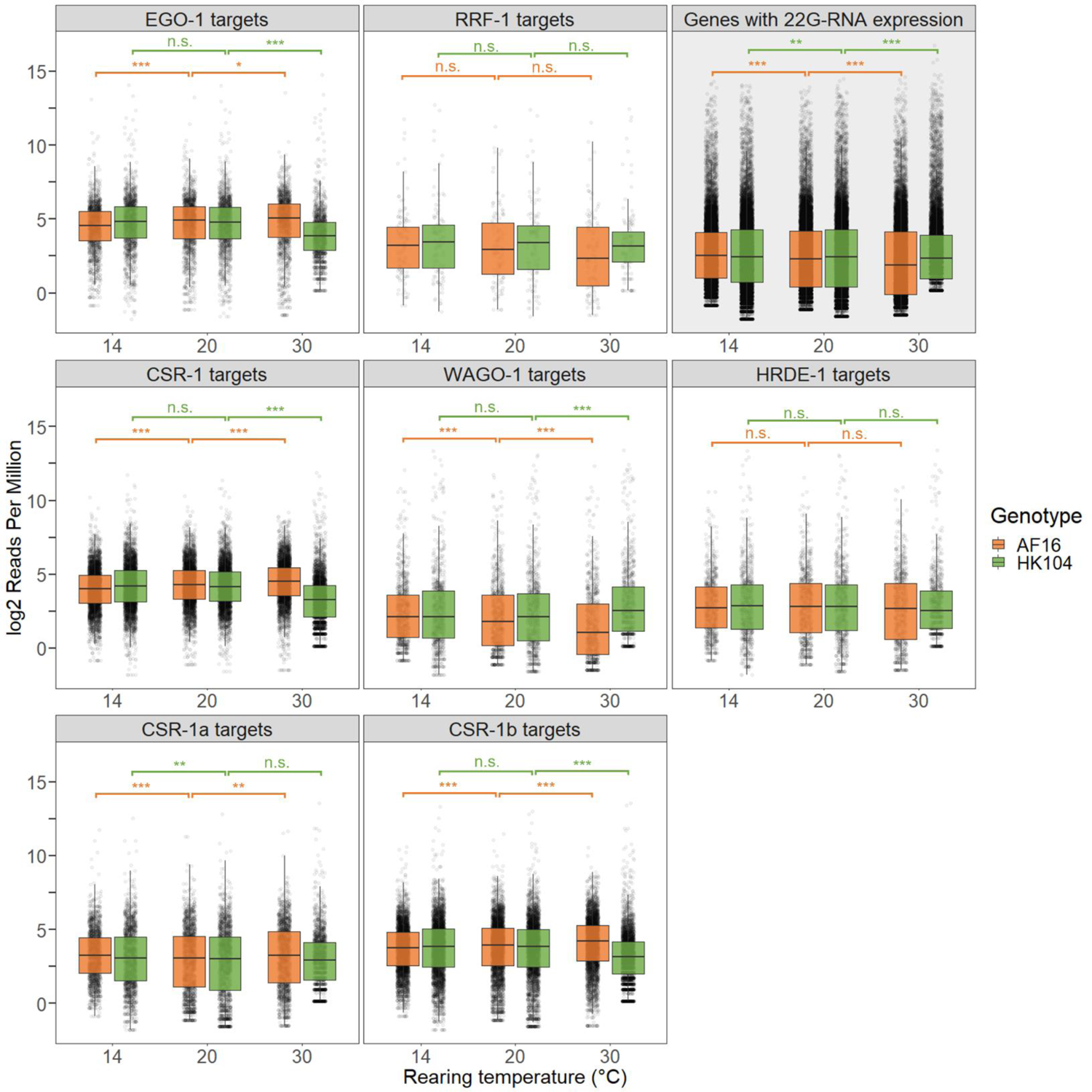
22G-RNA expression of genes targeted by EGO-1, RRF-1, CSR-1 (including both CSR-1a and CSR-1b), WAGO-1, or HRDE-1, as well as all genes showing 22G-RNA expression, for each genotype-temperature combination. 22G-RNA expression values were normalized using the TMM method to meet the assumption that most genes in the genome do not show differential 22G-RNA expression, followed by voom normalization to units of log_2_ RPM. Significant differences between temperature treatments of the same genotype were determined using a Wilcoxon rank sum test, with * indicating *p* < 0.05, ** indicating *p* < 0.01, *** indicating *p* < 0.001, and n.s. indicating non-significance.

**Table S1:** Proportion of 100kb bins along chromosome arm domains where mean 22G-RNA expression was lower at 30°C than it was at 20°C, for both genotypes. Bins showing decreased expression at 30°C were used to calculate the mean magnitude of the expression decrease for both genotypes. Expression decreases are in units of log_2_ Reads Per Million.

| **Chromosome** | **% of bins in AF16**<br>**with decreased**<br>**expression at 30 °C** | **% of bins in HK104**<br>**with decreased**<br>**expression at 30 °C** | **Average expression**<br>**decrease for AF16**<br>**bins (log<sub>2</sub> RPM)** | **Average expression**<br>**decrease for HK104**<br>**bins (log<sub>2</sub> RPM)** |
| --- | --- | --- | --- | --- |
| I | 64% (58 / 90) | 84% (76 / 90) | 0.55 | 1.33 |
| II | 70% (68 / 96) | 88% (84 / 96) | 0.72 | 1.60 |
| III | 62% (53 / 86) | 86% (74 / 86) | 0.57 | 1.33 |
| IV | 71% (67 / 95) | 80% (76 / 95) | 0.73 | 1.35 |
| V | 69% (80 / 116) | 77% (89 / 116) | 0.63 | 1.25 |
| X | 76% (112 / 147) | 82% (120 / 147) | 0.74 | 1.82 |

**Table S2:** Proportion of 100kb bins along chromosome center domains where mean 22G-RNA expression was lower at 30°C than it was at 20°C, for both genotypes. Bins showing decreased expression at 30°C were used to calculate the mean magnitude of the expression decrease for both genotypes. Expression decreases are in units of log_2_ Reads Per Million.

| Chromosome | % of bins in AF16<br>with decreased<br>expression at 30 °C | % of bins in HK104<br>with decreased<br>expression at 30 °C | Average expression<br>decrease for AF16<br>bins (log<sub>2</sub> RPM) | Average expression<br>decrease for HK104<br>bins (log<sub>2</sub> RPM) |
| --- | --- | --- | --- | --- |
| I | 64% (42 / 66) | 92% (61 / 66) | 0.46 | 1.49 |
| II | 55% (39 / 71) | 93% (66 / 71) | 0.42 | 1.43 |
| III | 39% (24 / 61) | 90% (55 / 61) | 0.31 | 1.42 |
| IV | 57% (46 / 81) | 85% (69 / 81) | 0.45 | 1.34 |
| V | 54% (43 / 80) | 89% (71 / 80) | 0.43 | 1.09 |
| X | 81% (57 / 70) | 91% (64 / 70) | 0.74 | 1.89 |

**Table S3:** voom-normalized expression of 22G-RNAs aligning antisense to 14,283 genomic features in all 17 replicates, in units of log_2_ RPM. “Gene_Biotype” indicates the type of feature (protein-coding gene, pseudogene, repeat, or transposable element). AF and HK refer to the AF16 and HK104 strains, respectively. 14, 20, and 30 indicate the rearing temperature of each replicate. “Differential_Expression_Category” indicates which category of differential expression each feature was assigned to.

**Table S4:** Targets of 22G-RNA pathway proteins in *C. briggsae* used in this study. CSR-1 targets are known for *C. briggsae*; targets of all other proteins are orthologs of known targets in *C. elegans*. Genes with available 22G-RNA or mRNA expression data are also indicated. “TRUE” indicates that a gene belongs to a given set of genes.

**Table S5.** Number of reads present in each replicate at various stages of the alignment and counting pipeline. AF and HK refer to the AF16 and HK104 strains, respectively. 14, 20, and 30 indicate the rearing temperature of each replicate. Statistics are for alignments to the respective genome (AF samples to the AF16 genome, and HK samples to the HK104 genome).

| **Replicate** | **Total # of raw reads** | **# of reads after trimming** | **# of reads aligned to the genome** | **# of reads aligned perfectly to the genome** | **# of perfectly-aligned reads aligned to an annotated feature** |
| --- | --- | --- | --- | --- | --- |
| AF14-1 | 9,714,767 | 8,624,894 | 7,596,951 | 6,664,095 | 4,859,004 |
| AF14-2 | 10,057,606 | 9,369,252 | 8,663,343 | 7,419,685 | 5,168,159 |
| AF14-3 | 9,153,959 | 8,341,891 | 7,837,217 | 6,934,035 | 5,704,460 |
| AF20-1 | 8,311,109 | 7,021,982 | 6,613,825 | 5,912,723 | 4,580,474 |
| AF20-2 | 7,187,732 | 6,047,269 | 5,738,135 | 5,153,967 | 3,975,678 |
| AF20-3 | 9,901,647 | 8,922,678 | 8,122,788 | 7,109,900 | 5,201,159 |
| AF30-1 | 21,556,148 | 20,809,661 | 18,949,949 | 16,009,149 | 11,612,300 |
| AF30-2 | 5,557,795 | 4,641,808 | 4,317,344 | 3,815,140 | 2,857,070 |
| AF30-3 | 13,334,748 | 11,638,513 | 10,736,202 | 9,141,443 | 7,362,039 |
| HK14-1 | 12,155,046 | 8,868,287 | 7,663,612 | 6,503,096 | 4,340,425 |
| HK14-2 | 11,165,762 | 9,530,022 | 8,239,591 | 6,958,250 | 4,565,104 |
| HK14-3 | 10,256,635 | 9,393,639 | 8,205,938 | 7,011,489 | 4,619,031 |
| HK20-1 | 11,667,443 | 10,624,773 | 9,621,489 | 8,498,648 | 5,800,746 |
| HK20-2 | 13,321,345 | 12,516,817 | 11,097,900 | 9,592,262 | 6,340,184 |
| HK20-3 | 11,495,618 | 10,514,634 | 9,462,109 | 8,163,107 | 5,842,908 |
| HK30-1 | 5,928,373 | 1,298,689 | 1,016,104 | 888,207 | 548,293 |
| HK30-2 | 19,716,033 | 15,924,990 | 13,997,848 | 12,362,971 | 7,941,578 |
| HK30-3 | 12,047,216 | 9,796,474 | 7,629,207 | 6,631,820 | 3,988,841 |

**Table S6.** Alignment statistics for all six genotype-temperature combinations. Small RNAs from each replicate were aligned to both the AF16 reference genome and HK104 pseudo-reference genome. Statistics for each replicate were averaged out within each genotype-temperature combination, excluding the outlier HK30-1 replicate.

| **Condition** | **Proportion of reads**<br>**aligning uniquely to**<br>**AF16 genome** | **Proportion of reads**<br>**aligning uniquely to**<br>**HK104 genome** | **Mismatch rate**<br>**per base against**<br>**AF16 genome** | **Mismatch rate**<br>**per base against**<br>**HK104 genome** |
| --- | --- | --- | --- | --- |
| AF16, 14°C | 74.0% | 72.7% | 0.22% | 0.34% |
| AF16, 20°C | 73.9% | 71.5% | 0.17% | 0.34% |
| AF16, 30°C | 70.7% | 69.1% | 0.21% | 0.34% |
| HK104, 14°C | 65.6% | 67.2% | 0.43% | 0.35% |
| HK104, 20°C | 68.8% | 70.4% | 0.36% | 0.29% |
| HK104, 30°C | 67.8% | 68.9% | 0.25% | 0.20% |

